# Mammalian retromer is an adaptable scaffold for cargo sorting from endosomes

**DOI:** 10.1101/639575

**Authors:** Amy K. Kendall, Boyang Xie, Peng Xu, Jue Wang, Rodger Burcham, Meredith N. Frazier, Elad Binshtein, Hui Wei, Todd R. Graham, Terunaga Nakagawa, Lauren P. Jackson

## Abstract

In metazoans, retromer (VPS26/VPS35/VPS29) associates with sorting nexin (SNX) proteins to form coats on endosomal tubules and sort cargo proteins to the *trans*-Golgi network (TGN) or plasma membrane. This core complex is highly conserved from yeast to humans, but molecular mechanisms of metazoan retromer assembly remain undefined. Here we combine single particle cryo-electron microscopy with biophysical methods to uncover multiple oligomer structures formed by mammalian retromer. Two-dimensional class averages in ice reveal the retromer heterotrimer; dimers of trimers; tetramers of trimers; and flat chains. These species are further supported by biophysical studies in solution. We provide cryo-EM reconstructions of all species, including pseudo-atomic resolution detail for key sub-structures. Multi-body refinement demonstrates how retromer heterotrimers and dimers adopt a range of conformations. Our structures identify a flexible yet highly conserved electrostatic interface in dimers formed by interactions between VPS35 subunits. We generate a structure-based mutant to disrupt this key interface *in vitro* and introduce equivalent mutations into *S. cerevisiae* to demonstrate the mutant exhibits a cargo sorting defect. Together, structures and complementary functional data in budding yeast imply a conserved assembly interface across eukaryotes. These data further suggest mammalian retromer acts as an adaptable and plastic scaffold that accommodates interactions with different SNXs to sort multiple cargoes from endosomes their final destinations.

## Introduction

Retromer is a multi-subunit protein complex that forms coats on tubules emerging from endosomes (reviewed in^1, 2^). The core heterotrimer, formerly called cargo-selective complex, contains VPS26, VPS35, and VPS29 subunits that together form a ~150kDa heterotrimer; for clarity, we will refer to the VPS26/35/29 heterotrimer as “retromer” in this manuscript. Retromer associates with sorting nexin (SNX) proteins that help recruit it to endosomal membranes enriched in phosphatidylinositol-3- phosphate (PI3P). In yeast, retromer is an obligate pentamer. Vps5 and Vps17 are SNX-BAR proteins that associate to form heterodimers^3^, and retromer/Vps5/Vps17 coats sort acid hydrolase receptors, including Vps10, from endosomes to the *trans*-Golgi network (TGN)^3^. In metazoans, retromer interacts with multiple sorting nexin proteins (SNX-BARs, SNX3, and SNX27) to diversify its cargo repertoire. Hydrolase receptors including mannose 6-phosphate receptors^4^ are sorted in a retrograde pathway to the TGN, while other transmembrane receptor cargoes (e.g. β2-adrenergic receptor, GLUT1) are recycled directly from endosomes to the plasma membrane^5, 6^. Mammalian retromer uses additional SNX proteins, including SNX3^7^ and SNX27^8, 9^, as cargo adaptors. SNX3/retromer is implicated in Wntless retrograde sorting^10–14^, while SNX27 sorts many transmembrane cargoes bearing specific PDZ binding motifs to the plasma membrane^5^. SNX-BAR proteins have more recently emerged as cargo adaptors^15, 16^, and VPS26 itself directly binds at least one cargo^17^.

Mechanisms governing the assembly and structures of coated tubules are now starting to emerge (recently reviewed in ^18, 19^). Multiple crystal structures of mammalian retromer subunits ^20–23^ and partial sub-complexes ^24, 25^ have been determined. Two groups have proposed mammalian retromer forms a “dimer of trimers” ^25, 26^ based on low resolution small-angle X-ray scattering data, and a structure has been proposed for the budding yeast heterotrimer from electron microscopy (EM) data ^27^. A recent cryo-electron tomography (cryo-ET) structure reveals the overall architecture of reconstituted thermophilic yeast retromer with a Vps5 homodimer ^28^. Whether metazoan retromer assembles in the same way remains an open question. Known cargo- or PI3P-binding sites for SNX3 and SNX27 seem to be located far from the membrane^18, 28^ when partial crystal structures^8, 25^ are docked into the yeast cryo-ET model. SNX proteins lacking BAR domains may form different retromer assemblies to sort distinct cargoes.

Here we present structural and biophysical data to reveal how mammalian retromer forms a variety of oligomers. We find retromer forms dimers of trimers; a tetramer of trimers; and longer chains. Our structures suggest retromer assembles via a key electrostatic yet flexible interface mediated by the backside of VPS35 subunits. We test this proposed interface biochemically by introducing structure-based point mutations and determining whether retromer can form oligomers *in vitro*. Finally, we introduce equivalent mutations into *S. cerevisiae* and show that disrupting the conserved interface impedes cargo sorting. Disruption of this interface with specific point mutations suggests the interface may be relevant for assembling higher order structures *in vivo*. Based on our structures, retromer appears to function as a plastic and adaptable scaffold, and we propose possible assembly models for the scaffold in the presence of sorting nexins on endosomal membranes.

## Results

### Structural & biophysical studies of mammalian retromer

In preliminary studies, we analyzed purified recombinant retromer suitability for single particle cryo-EM studies. We first obtained two-dimensional (2D) class averages and produced random conical tilt reconstructions from negatively stained samples (data not shown). 2D class averages from untilted particles revealed heterotrimers along with multiple oligomeric species, including dimers and tetramers of trimers (data not shown). We next undertook single particle cryo-EM studies (Figure 1) to determine structures of mammalian retromer heterotrimer (Figure 1A) and oligomers (Figures 1B-E). In vitrified ice, we clearly observe the retromer heterotrimer; dimers of trimers; and tetramers of trimers both in micrographs (Figure S1) and in 2D class averages (Figure 1). We also observe longer, flat chains of retromer (Figure 1C, 1D). We used 2D classification to separate each biochemical species and generated reconstructions for each (workflow in Figure S2; full details in Materials & Methods and Extended EM Methods). We independently confirmed retromer forms oligomers in solution using size exclusion chromatography with multi-angle laser light scattering (SEC-MALS; Figure S3A) and dynamic light scattering (Figure S3B). We find oligomer formation depends on salt concentration; the retromer heterotrimer forms both dimers and tetramers near physiological salt concentrations (50-100 mM NaCl) but exists as a heterotrimer at higher concentrations (150-500 mM NaCl). We describe each structure and its key features below.

**Figure 1.**
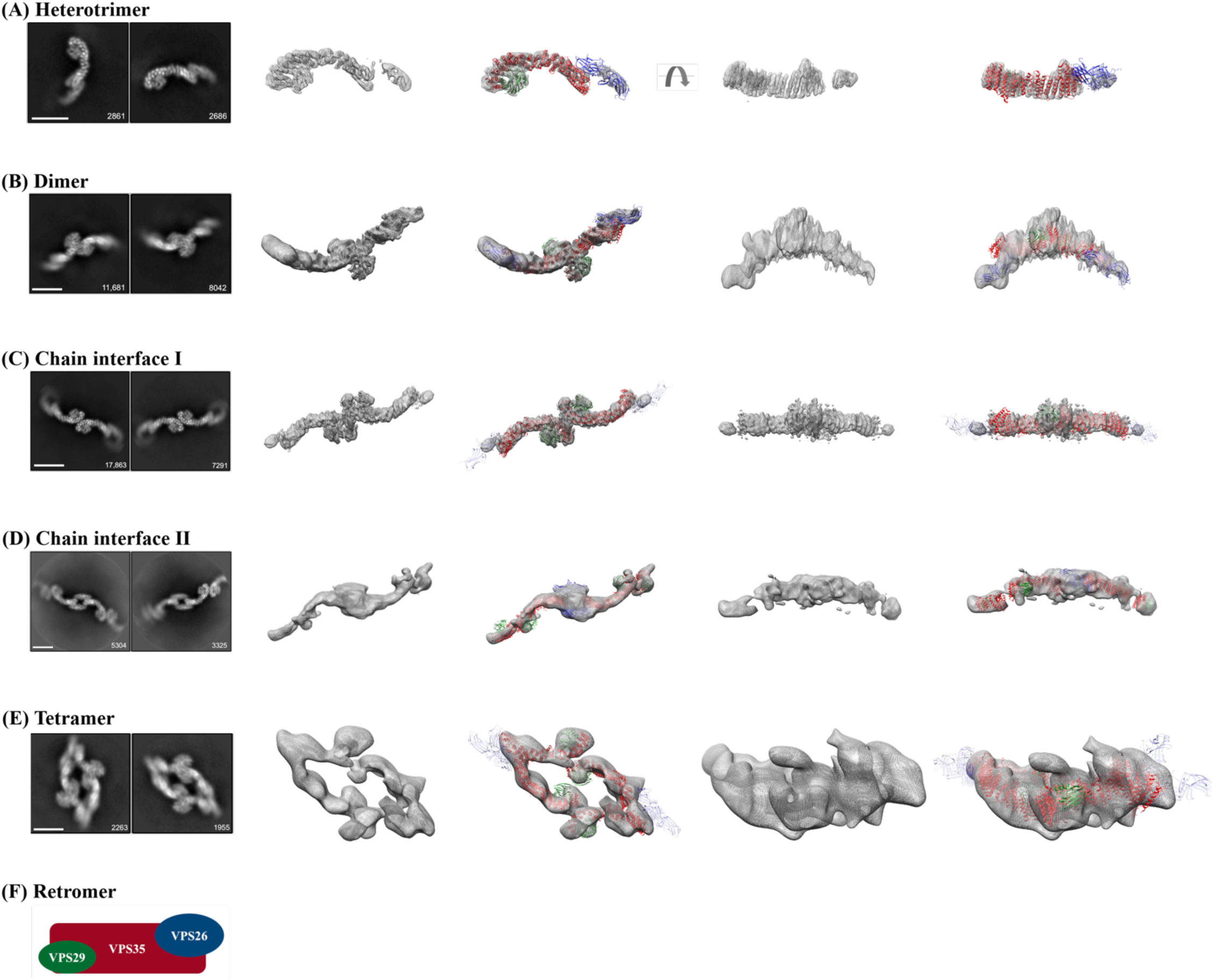
Single particle cryo-EM reconstructions of mammalian retromer. Four retromer species were resolved between 27 and 6 Å: (A) the retromer heterotrimer; (B) a dimer of trimers; (C, D) retromer chains centered on two different interfaces; and (E) a tetramer of trimers. For each row (A-E), the first column shows two representative 2D class averages for the species, including particle numbers. Scale bars represent 10 nm. The middle two columns show 3D reconstructions (see Figure S2 for contour details) with and without a fitted model. The last two columns show an additional view (rotated by 90°) of 3D reconstructions with and without fitted models. Initial models were generated from partial crystal structures (PDB ID: 2R17, 5F0J). VPS29 is shown in green, VPS35 in red, and VPS26 in blue or transparent blue (when averaged out in a reconstruction). A schematic of the retromer heterotrimer is shown in (F).

#### Heterotrimer

The structure of full-length mammalian retromer is directly observed here for the first time (6.0 Å average resolution from 26,369 particles). 3D reconstructions reveal a well-resolved interface between the VPS35 C-terminus (C-VPS35) and VPS29. This C-VPS35/ VPS29 interface is nearly identical to a partial X-ray crystal structure (PDB: 2R17), which we used as a model for fitting into reconstructions. The resolution of the map corresponding to C-VPS35/VPS29 is 5.5 Å (Figure S4A), and α-helical density for the VPS35 solenoid is clearly distinguishable and can be fit well (Figure S4B, S4C). In contrast, the VPS35 N-terminus exhibits substantial flexibility. The VPS35 N-terminal interface that binds VPS26 is not well-ordered, and this part of the map is less-ordered (~8-9 Å; Figure S4A). VPS26 is likely not well-resolved for two reasons. First, VPS26 is an all β-sheet protein, and sheets will not be as well-resolved in this resolution range. Second, both N-VPS35 and VPS26 appear to be flexible (discussed further below).

#### Retromer dimer of trimers

Retromer forms dimers of trimers in both negatively stained (data not shown) and vitrified samples (Figure 1B). The dimeric interface between two heterotrimers is immediately clear in 2D class averages, because we can distinguish between the VPS35 and VPS26 ends of retromer. Reconstructions (14.6 Å average resolution from 31,022 particles) reveal the C-termini of VPS35 subunits mediate dimer formation. The map is slightly better resolved at the dimer interface mediated by VPS35 (14 Å resolution; Figure S4D). The mammalian retromer dimer is wide, fairly flat, and does not exhibit two-fold symmetry. Instead, one heterotrimer twists and rotates relative to its partner, giving rise to a dimer with an angle of ~150° between two heterotrimer legs. The retromer dimer is a stable biochemical species (Figure S3), and like the heterotrimer, it exhibits substantial conformational flexibility (see next section). The VPS35-mediated interface is slightly different across the population of dimers, giving rise to slightly different dimers with different angles between heterotrimer legs. Furthermore, the presence of VPS26 seems to affect dimer formation and overall retromer structure. Class averages of negatively stained VPS35/VPS29 sub-complex (Figure S6A) reveal dimers that are more flexible than dimers formed by intact retromer. VPS26 appears to impart rigidity to the VPS35 solenoid, even though structural data from heterotrimers and dimers reveal flexibility in the N-terminal VPS35/VPS26 interface. Overall, these data suggest retromer is plastic and adaptable at multiple interfaces.

#### Multibody refinement

2D classes and 3D reconstructions suggest both the heterotrimer and dimers exhibit substantial structural heterogeneity. We thus implemented multi-body refinement in RELION-3^29^ to model different portions of retromer as rigid bodies. We chose three rigid body units for the refinement: VPS26; N-VPS35 (helices 1-15; residues 1-365); and C-VPS35 (helices 16-33; residues 365-780)/VPS29. The heterotrimer thus contained three rigid bodies, while dimers contained six. We established contributions of all eigenvectors to the total variance in both the heterotrimer (Figure 2A) and in dimer (Figure 2F) populations. The top three eigenvectors in the heterotrimer (Figure 2A) represent 49% of the data. We captured the range of heterotrimer conformations observed in the data (Figure 2B-D; Movie S1, S2). We observe flexibility in the middle of the VPS35 solenoid between helices 14 and 15; this implies retromer contains a “hinge point” to allow the N- and C-terminal halves of the VPS35 solenoid to move relative to each other (Movie S1, S2). These structural data, captured directly from a population of individual particles for the first time, suggest the heterotrimer itself exhibits inherent plasticity. Dimers are extremely heterogeneous, as the top three eigenvectors represent only 33% of the data (Figure 2F). Both visual representations (Figure 2G-I) and Movie S2 demonstrate heterotrimers move with respect to each other within the dimer. Overall, this flexibility likely has important biological consequences (see Discussion), which could allow retromer to accommodate multiple sorting nexins^8, 28^ and cargoes ^17^.

**Figure 2.**
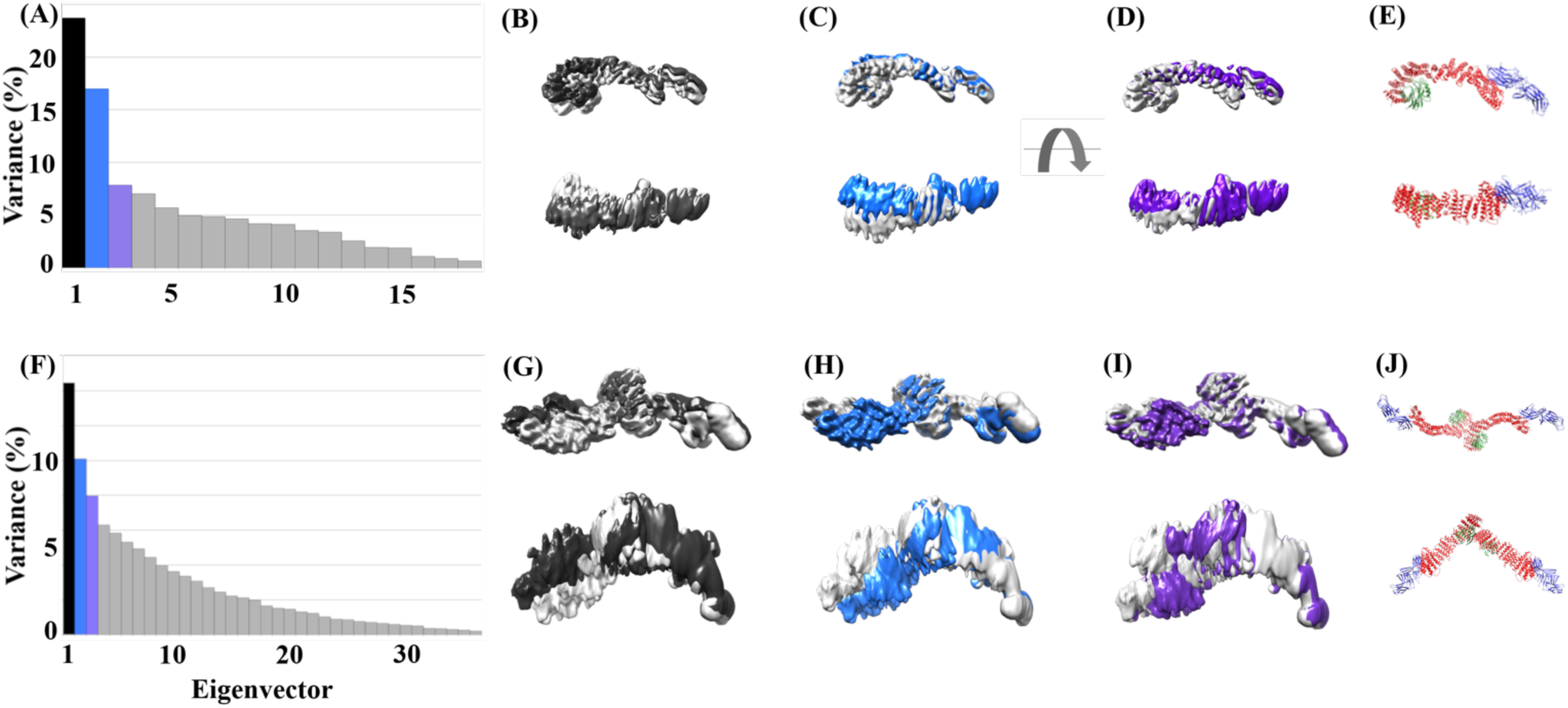
Retromer particles exhibit conformational flexibility. The retromer heterotrimer and dimer underwent multi-body refinement in RELION to characterize structural heterogeneity and flexibility. Contributions of all eigenvectors to heterotrimer variance are shown in (A). (B-D) show two different views (top & bottom, rotated by 90°) of heterotrimer movements represented by the top three eigenvectors; colors match graph shown in (A). The VPS35 subunit possesses a “hinge point” that allows its N- and C-termini to move with respect to each other. Contributions of all eigenvectors to dimer variance are shown in (F). (G-I) show two different views (top & bottom, rotated by 90°) of dimer movements represented by the top three eigenvectors. The VPS35/VPS35 dimer interface is flexible, which leads to large movements in one heterotrimer with respect to its partner. (E) and (J) show equivalent heterotrimer and dimer ribbon diagrams for orientation. VPS29 is shown in green, VPS35 in red, and VPS26 in blue.

#### Retromer chains

A surprising result from our structural studies was the existence of longer retromer chains in vitrified ice (Figure S1C, S1D; Figure 1C, 1D). We did not observe chain structures in negative stain, and they may occur only at higher concentrations present in vitrified samples and when retromer is concentrated on membranes. Chain structures arise when retromer links together at both the C-VPS35 (Figure 1C) and VPS26 ends (Figure 1D); VPS26-mediated interfaces resemble a chain link. We observed chains as short as 3 links (Figure S1C) and up to ~20 links, because chains can span an entire micrograph (data not shown). To analyze the structure of chains computationally, we used masks to separate chains into two different interfaces: the first interface was centered on VPS35-mediated dimers (Chain interface I; Figure 1C), while the second was centered on VPS26 chain links (Chain interface II; Figure 1D). We specifically set out to compare the VPS35-mediated dimer found in chains with the curved VPS35 interface observed in dimers (discussed above). We generated 3D reconstructions of the VPS35 dimer from chains (7.1 Å average resolution from 75,790 particles), which is well-resolved at the dimer interface (Figure S4E). This VPS35-mediated interface appears flat and exhibits 2-fold symmetry.

We also generated reconstructions of the VPS26-mediated interface (Figure 1D; Figure S4G) at 18.5 Å average resolution from 13,782 particles. Retromer chains exhibit preferred orientation (Figure S2), which limits the resolution of our model. The data do not allow us to build an accurate pseudo-atomic model that captures how VPS26 subunits pack together. However, we clearly see a symmetrical two-fold interface mediated by VPS26 in 2D classes. These links allow adjacent VPS35 subunits to curve out and away from each other, and we clearly observe the curve of VPS35 solenoids in both 2D classes and 3D reconstructions. Alternating heterotrimer units in this way would allow retromer to form an elongated repeating structure. Overall, the existence of retromer chains suggests the heterotrimer alone may encode assembly information to build a flexible scaffold to accommodate various binding partners.

#### Retromer tetramer of trimers

A second surprising result from EM studies was the presence of stable biochemical tetramers (Figure 1E). We observe tetramers in both micrographs and 2D class averages obtained from negatively stained (data not shown) or vitrified retromer (Figure S1B). Their presence is further supported by biophysical data (Figure S3). Tetramers are the rarest species in our samples (6,015 particles) but the overall architecture is relatively clear. Tetramers assemble using two interfaces: the first is a curved VPS35-mediated dimer interface similar to that observed in the dimer species. The second interface is mediated by VPS26, although this interface is very poorly resolved. Most of our views are “top down” views, and limited side views suggest a curved particle shaped like a shallow boat or bowl. However, the low particle numbers; inherent VPS26 flexibility; and limited views impede our ability to generate reconstructions beyond 27 Å (Figure 1E; Figure S4G).

### VPS35 contains a conserved electrostatic interface to mediate dimer assembly

We observed two different dimers mediated by C-VPS35 subunits in our retromer oligomers (Figure 1B, 1C); one dimer is flat, while the second appears curved. Dimer formation seems to be an inherent property of VPS35, since dimers form in solution (Figure S3) and on grids in the absence of sorting nexins or other binding partners. We tested whether C-VPS35 plays a role in promoting higher-order retromer assembly. VPS35 α-solenoid regions consistently produced the best-resolved portions of 3D reconstructions (Figure S4A, S4B, S4E). We therefore pursued sub-structures of VPS35-mediated interfaces observed in retromer dimers and in chain interface I (Figure 3; Figure S2). In both species, we masked the VPS35 N-terminus and VPS26 subunits in order to focus on C-VPS35/VPS29 (Figure 3A, 3B). We produced reconstructions of each dimer sub-structure (Figure 3C, 3D).

**Figure 3.**
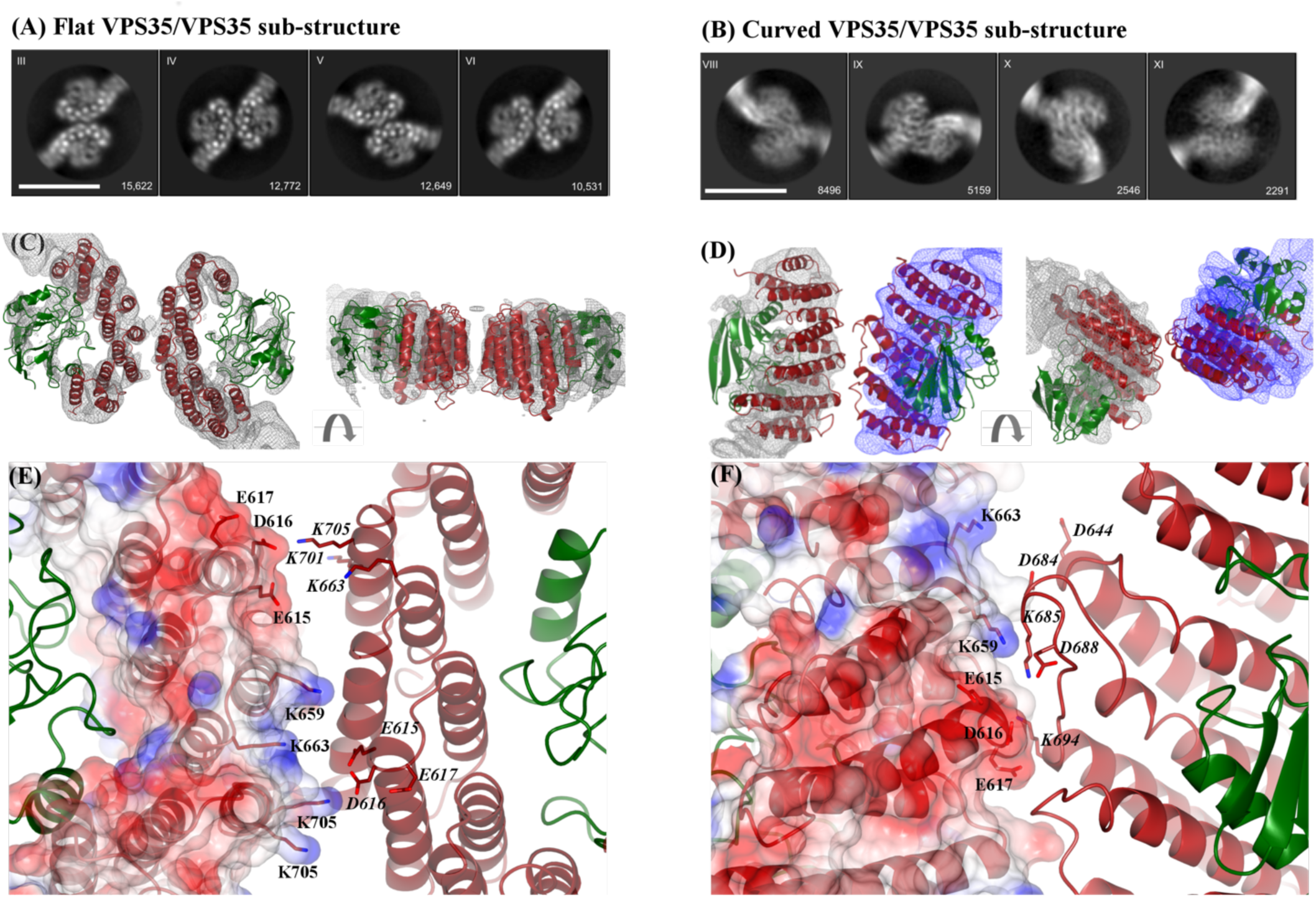
Conserved electrostatic patches mediate key VPS35 dimer interfaces. Representative 2D class averages of the (A) VPS35/VPS35 flat dimer sub-structure generated from retromer chain interface I and (B) the VPS35/VPS35 curved sub-structure generated from retromer dimers. Scale bars represent 10 nm. (C) Two different views (rotated 90°) of flat sub-structure reconstruction at 5.3 Å shown as grey mesh (contoured at 4σ) with fitted model overlaid (PDB: 2R17). VPS35 is shown in red, VPS29 in green. (D) Two different views of curved sub-structure reconstruction at ~6 Å with fitted model. Reconstructions are shown as grey and blue mesh (contoured at 4σ); each partner was processed separately in multi-body refinement (details in text.) (E) Inset shows detailed view of the flat dimer interface, which exhibits two-fold symmetry. One VPS35 copy is shown as an electrostatic surface with residues marked in black text; the second copy is shown as a ribbon diagram with the same residues shown in italics. Key acidic residues in one copy (E615, D616, E617) interact with basic residues in the second copy (K659, K663, K701, K705). (F) Inset shows detailed view of curved dimer interface, which does not exhibit two-fold symmetry. The interface is shown as described in (E). Many of the same residues are observed in both dimer interfaces, including E615, D616, E617, K659, and K663.

The sub-structure reconstruction from chain interface I (Figure 3A, 3C; 69,195 particles at 5.0 Å average resolution) revealed a symmetrical and flat two-fold interface. In contrast, the sub-structure reconstruction from dimers (Figure 3B, 3D; 32,435 particles at 5.5 Å average resolution) revealed a curved orientation between VPS35 subunits, which does not exhibit two-fold symmetry. We clearly observe α-helices in 2D classes (Figure 3A) and 3D reconstructions (Figure 3B) of flat dimers. The curved sub-structure was more challenging because of inherent flexibility; this sub-structure required us to separate and process the two bodies using multi-body refinement (cf. Figure 2F-I).

Overall, improved reconstructions from sub-structures allowed us to generate pseudo-atomic models for each dimer using the C-VPS35/VPS29 partial crystal structure (PDB: 2R17) as an initial model. Both models reveal dimer interfaces located on the back side of VPS35 subunits, on the opposite face from where VPS35 binds VPS29 (Figure 3C, 3D). The flat dimer exhibits 2-fold symmetry, while the curved dimer shows one C-VPS35/VP29 has rotated away from its partner. In both dimers, VPS29 subunits point out and away from VPS35. VPS29 residues known to mediate interactions with key regulators, including VARP ^30^ and TBC1D5 ^31, 32^, are fully exposed in each dimer. Maps from the flat sub-structure were well-resolved (Figure 3C), because the structure is symmetrical and is not flexible, so we refined the model using real space refinement in PHENIX (Table S2; Supplemental methods). Maps from the curved sub-structure are poorer due to high flexibility and lack of symmetry, so we could not justify further refinement. However, reconstructions of the curved sub-structure allow us to observe overall arrangement of secondary and tertiary structural elements in order to compare the two dimer interfaces.

Both dimer interfaces contain multiple electrostatic residues (Figure 3E, 3F). The structures are further supported by our biophysical data, which show retromer oligomer formation depends on salt concentration (Figure S3). Many of the same residues are observed in both interfaces, including E615, D616, E617, K659, and K653. In the flat dimer, acidic residues (E615, D616, E617) in one copy associate with basic residues (K663, K701, K703) in its symmetry copy (Figure 3E). In the curved dimer, we observe the same acidic residues (E615, D616, E617), but they mediate different contacts with K685 and K694. The resolution of our sub-structures is insufficient to measure distances accurately. However, the pseudo-atomic model of the flat sub-structure suggests these electrostatic residues are located within 7-8 Å, which would be close enough to mediate salt bridge formation. Many key residues in both interfaces are highly or absolutely conserved from yeast to humans (Figure S5A), suggesting they play an important biological role. We compared our interfaces with other proposed retromer dimers. Our interfaces differ from mammalian dimers proposed from SAXS data ^25, 26^, which suggest an end-to-end VPS35 dimer. Our structures are more similar to a recent model proposed from *C. thermophilum* retromer ^28^ assembled on membranes *in vitro*. The cryo-ET model cannot resolve side chains, but our proposed residues are both conserved in *C. thermophilum* and are located in the VPS35 dimer interface proposed to form arch-like structures (Figure S5B). Based on available sequence and structural data, these electrostatic residues make excellent candidates for testing structure-based models (next section).

### Mutating the conserved interface disrupts assembly and impedes cargo sorting

We next tested whether key residues in our proposed VPS35/VPS35 interfaces would promote assembly. Briefly, we introduced point mutations in VPS35 subunits predicted to break apart the electrostatic interface and generated recombinant purified retromer protein containing mutant VPS35 subunits with wild-type VPS26 and VPS29. To test assembly *in vitro*, we combined size exclusion chromatography with negative stain electron microscopy to test whether point mutations affect whether retromer forms higher order oligomer structures. We identified five candidate residues conserved from yeast to humans (Figure S5A): E615, E617, K659, K662, and K663. We also included D616, which is conserved from insects to vertebrates. Initially, we attempted a charge swap in which we mutated three conserved lysine residues (K659, K662, K663) to glutamates. We predicted this should prevent dimer formation by causing VPS35 subunits to repel each other. This mutant gave a partial phenotype (Figure S6B); we occasionally observed dimers in negatively stained samples but at much lower frequency than in wild-type samples (data not shown). We therefore introduced three additional mutations to generate the retromer electrostatic mutant, AAA3KE (E615A/D616A/E617A/ K659E/K662E/K663E). This mutant exhibited a clear shift in gel filtration profile (Figure 4A). The primary wild-type retromer peak elutes predominantly at volumes consistent with tetramers over gel filtration, while the electrostatic mutant elutes predominantly as a heterotrimer. This suggested we successfully mutated residues that mediate assembly. We could not identify tetramers or dimers in micrographs of negatively stained AAA3KE mutant samples (data not shown). However, representative 2D class averages of negatively stained mutant AAA3KE protein revealed stable heterotrimers (Figure 4B); this confirms we did not disrupt overall heterotrimer fold. Together, these data suggest we successfully disrupted a key interface mediating formation of oligomers formed by mammalian retromer.

**Figure 4.**
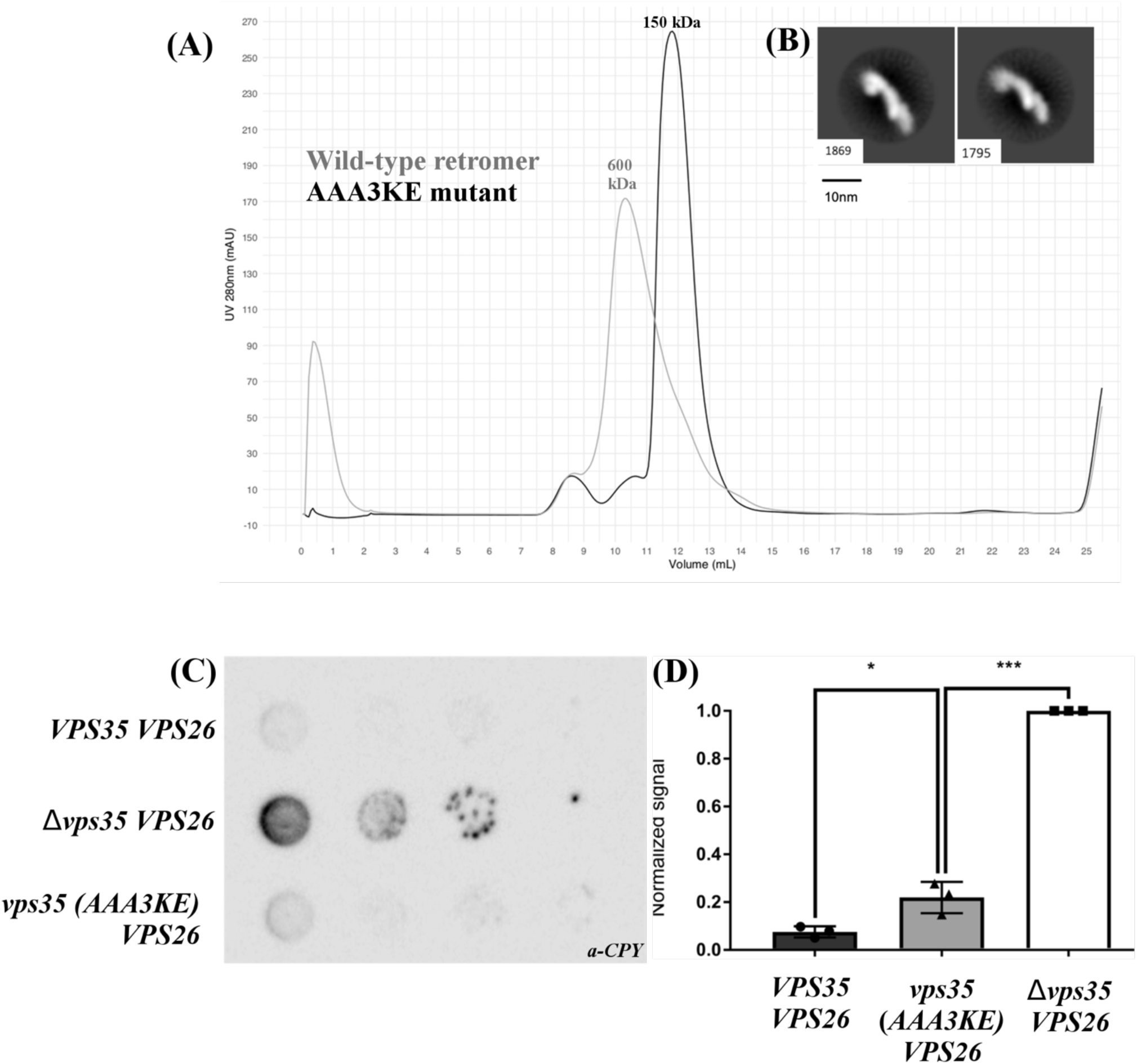
Mutating the conserved VPS35 interface disrupts assembly and impedes cargo sorting in budding yeast. (A) Gel filtration profiles of wild-type mammalian retromer (grey trace) and the electrostatic AAA3KE mutant (E615A/D616A/E617A/K659E/K662E/K663E; black trace). The main wild-type peak elutes at a volume consistent with larger oligomeric assemblies (~600kDa), while the electrostatic mutant peak elutes at a volume consistent with the heterotrimer (~150kDa). (B) Two representative 2D class averages of the electrostatic mutant in negative stain. The overall 3D structure of the heterotrimer is preserved. Scale bar represents 10 nm. (C) One representative blot from a CPY secretion experiment in the *ΔVPS35ΔVPS26* retromer strain at 26°C. (D) Quantification from three biological replicates for each genotype in CPY secretion experiments. Data were analyzed using a one-way ANOVA with Tukey-Kramer post-hoc test. CPY secretion was normalized to the *ΔVPS35* yeast strain (set at 1.0). The retromer electrostatic mutant secretes about twice asas much CPY compared to wild-type retromer. Raw Western blots from all three replicates are shown in Figure S7.

We next tested whether our mutant would disrupt retromer assembly *in vivo*. Reconstituted Vps5/retromer from thermophilic yeast assembles VPS35-mediated dimers in the presence of sorting nexins on membranes. Our dimers and chains do not exhibit the same architecture without SNXs (see Discussion for details), but all three interfaces contain these highly conserved electrostatic residues. We next undertook a functional cargo sorting assay in budding yeast, *S. cerevisiae*, to determine whether breaking the conserved assembly interface would have a functional consequence. We used the well-characterized carboxypeptidase Y (CPY) secretion assay ^33^. Briefly, budding yeast sort CPY to the vacuole when retromer is present and functional. Loss of retromer instead drives CPY secretion through mis-sorting of Vps10, and CPY secretion is measured and quantified at the cell surface using an immunoblotting assay.

We generated a *Δvps35Δvps26* deletion strain from a *Δvps35* strain (kindly provided by Scott Emr’s lab, Cornell). We then re-introduced either wild-type or mutant retromer *VPS35*, together with wild-type *VPS26*, on plasmids under control of endogenous promoters and measured how much CPY was secreted in cells containing wild-type or mutant retromer. We normalized our data against a strain lacking *VPS35* by arbitrarily setting its CPY secretion at 1.0. The recombinant wild-type retromer strain, in which both *VPS35* and *VPS26* genes were re-introduced, secreted CPY at ~10% of Δ*vps35* levels (Figure 4C, 4D). The *vps35* AAA3KE mutant secreted about twice as much (~20%) CPY as compared to wild-type, across three biological replicates.

Mutating key conserved residues in the VPS35 dimer interface thus causes a reproducible and measurable cargo sorting defect. However, loss of these residues does not preclude cargo sorting in budding yeast. This may reflect the ability of SNX proteins to sort cargoes like CPY, as SNX1/SNX2 does in mammalian systems ^16, 34^. Mutating the heterotrimer scaffold may partially destabilize the coat, but it is insufficient to block cargo sorting, at least in yeast. This may be partly explained by the adaptability of the retromer scaffold (discussed below).

## Discussion

We have shown here how the conserved retromer heterotrimer (VPS26/VPS35/VPS29 subunits) forms stable oligomers. Single particle cryo-EM reconstructions provide the first structural snapshots of multiple mammalian retromer oligomers, including heterotrimer and dimer sub-structures at pseudo-atomic resolution. Previous reports ^25, 26^ identified a mammalian dimer based on very low resolution SAXS data, in which two groups proposed dimers mediated by the very C-terminal tips of VPS35 subunits. Our pseudo-atomic resolution data instead support an interface mediated by key conserved electrostatic residues located on the backside of VPS35 C-termini. One of our dimer models is similar to a published thermophilic yeast cryo-ET model ^28^ (discussed further in next section). Both biochemical and complementary functional data in budding yeast suggest this conserved interface mediates retromer dimer assembly. Identification and verification of specific electrostatic residues in the interface provides a useful molecular tool to the community for testing retromer function more precisely with different cargoes across multiple model systems.

### Implications for retromer assembly & regulation

Our structural data suggest possible models for retromer assembly (Figure 5). We generated models for structures with sorting nexins based on the curved and flat VPS35-mediated dimer interfaces, because we obtained reconstructions and tested specific residues in these interfaces. We generated possible mammalian retromer structural models with two sorting nexins, SNX27 and SNX3, by overlaying partial crystal structures (PDB ID: 4P2A and 5F0J, respectively) onto our cryo-EM reconstructions (Figure 5). The considerable VPS26 flexibility in both heterotrimers (Figure 2A; Movie S1, S2) and dimers (Figure 2; Movie S3) make it challenging to depict the position and orientation of VPS26. The position of VPS26 in our data could be consistent with that observed in a partial crystal structure^25^ and cryo-ET reconstructions^28^. We have chosen to model it based on the crystal structure here, because these are the highest resolution data available. However, our data indicate substantial flexibility at the VPS26/N-VPS35 interface, so VPS26 may adopt multiple conformations.

**Figure 5.**
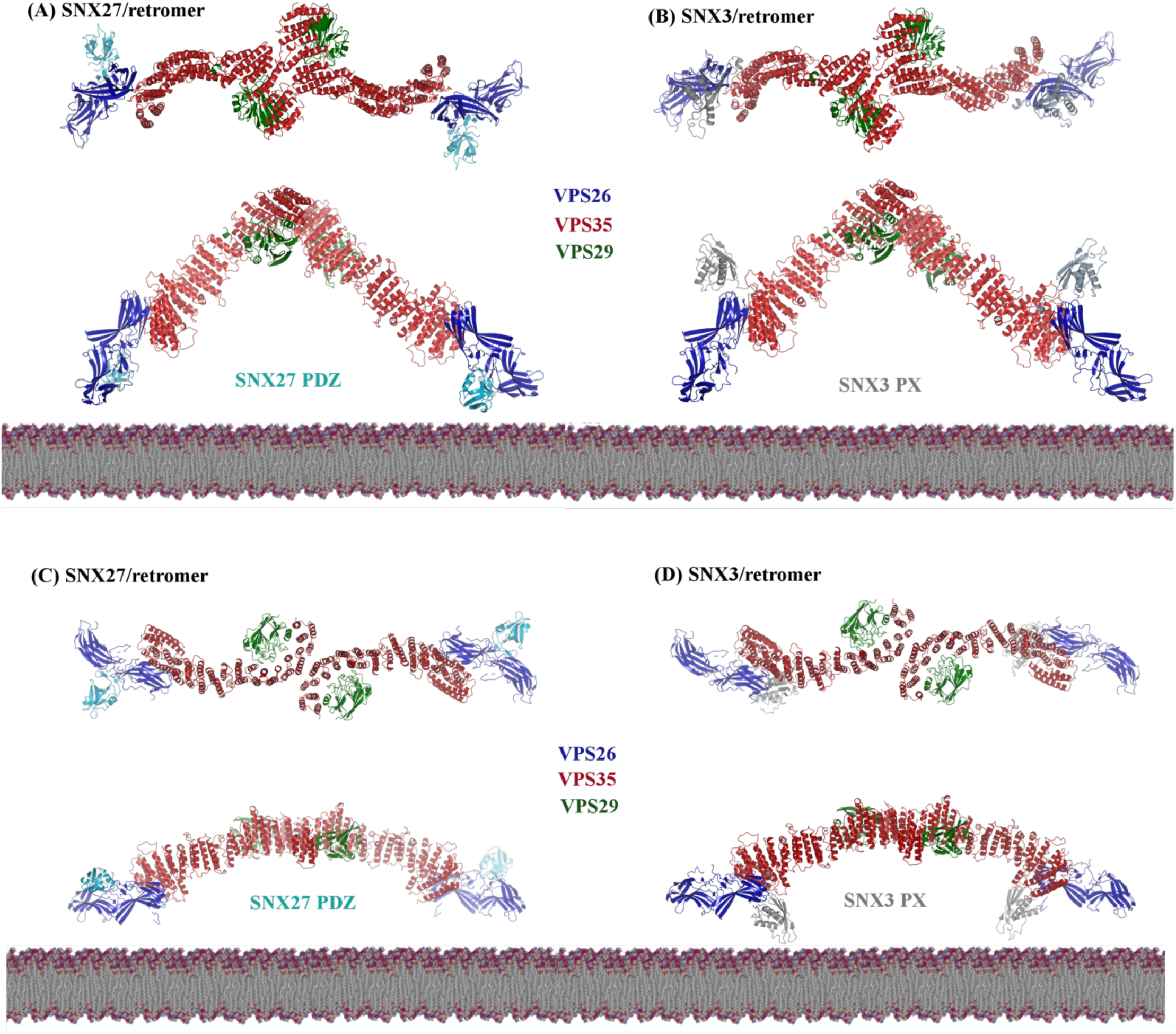
Retromer is a plastic scaffold for cargo sorting on membranes. Possible models of assembled retromer dimers (based on this work) shown with mammalian sorting nexins, SNX27 and SNX3. The left-hand column shows putative models of curved (A) or flat (C) retromer dimers with the SNX27 PDZ domain; models were generated by overlaying dimers with the VPS26/SNX27 PDZ crystal structure (PDB: 4P2A). Both retromer dimers place the SNX27 PDZ domain close to the membrane, where it could engage cargoes containing PDZ-binding motifs. The SNX27 PX and FERM domains are not shown, because there are no structures of full-length SNX27 to show overall architecture. The right-hand column shows models of curved (B) or flat (D) retromer dimers with the SNX3 PX domain; models were generated by overlaying dimers with the N-VPS35/VPS26/SNX3 PX crystal structure (PDB: 5F0J). Only flat retromer dimers (D) place the SNX3 PX domain close to the membrane in an orientation where known PX residues could interact with PI3P.

Models of curved, arch-like retromer dimers (Figure 5A, 5B) position the SNX27 N-terminal PDZ domain close to the membrane (Figure 5A). There are no structures of full-length SNX27, so we do not know how the PDZ domain sits relative to PX and FERM domains. It is possible retromer could sit in this orientation, but it remains unclear how retromer would then form a repeating unit. The SNX3/retromer model positions the SNX3 PX domain relatively far from the membrane (Figure 5B), and we think it unlikely retromer would assemble a coat in this way. Binding to either SNX27 or SNX3 may re-orient or lock a specific VPS26 conformation on a membrane, so that it becomes less flexible. Overall, extended arch-like dimers observed here seem more likely to be partial retromer assemblies in solution or the cytoplasm. We predict encountering specific SNXs or cargoes in PI3P-enriched membranes will correctly orient and position the retromer scaffold depending on biological need (i.e. what cargoes require sorting).

The existence of flat chains may suggest a new assembly mechanism for mammalian retromer. Formation of flat chains could position both SNX27 PDZ (Figure 5C) and SNX3 PX domains (Figure 5D) close to the membrane. SNX27/retromer coats would presumably be further oriented through binding PI3P via the PX domain and NPxY cargoes via the FERM domain. The PI3P-binding pocket for SNX3/retromer would also be apposed to the membrane. Chains further suggest a way for VPS26 subunits to form interfaces that extend retromer into repeating units (Figure S8). These chains would form very elongated structures with shallow and gently curving arches. These chains would position VPS29 subunits at arch apexes closer to the membrane, which in turn may have regulatory implications. Both VARP ^35^ and TBC1D5 ^31, 32^ interact with VPS29, and key VPS29 residues are exposed and available in all of our models. VARP contains two cysteine-rich repeats, and both have been shown to bind the same patch (L152) on VPS29 ^35^. Together with structural data, this implies one VARP could “bridge” a retromer VPS35-mediated dimer. VARP also binds endosomal Rab proteins and the R-SNARE, VAMP7. We can speculate that VARP may favor binding a flatter dimer located closer to the membrane, or alternatively, VARP could prompt a conformational change as part of its regulatory role.

### Comparison with yeast Vps5/retromer structure

The cryo-ET reconstruction of thermophilic yeast Vps5/retromer revealed how the retromer heterotrimer forms two types of dimers when assembled *in vitro* on membranes in the presence of a SNX-BAR protein ^28^. A symmetrical VPS26-mediated dimer is organized directly through interactions with a Vps5 homodimer. VPS35 subunits mediate formation of a second dimer that forms arches. We attempted to address two questions arising from the cryo-ET model in our work. Does the VPS35-mediated dimer exist only in the presence of SNXs, and is the dimer conserved across evolution? Our biochemical and structural data agree with previously published reports ^25, 26^ indicating mammalian retromer forms dimers on its own in solution. We have now visualized those dimers, as well as new chain and tetramer structures, here for the first time. Together, these data imply the heterotrimer acts as a scaffold with intrinsic assembly capabilities. We further show how VPS35 contains a specific electrostatic interface, because we can disrupt dimer assembly with targeted point mutations (Figure 4). We further tested for functional conservation in the interface using the CPY secretion assay. Our data suggest disrupting the retromer scaffold has a modest but reproducible effect on CPY secretion in budding yeast. CPY is sorted by Vps10, and in mammalian cells, the Vps10 homologue (CI-MPR) has been shown to depend upon sorting nexin dimers (SNX1/2) instead of the heterotrimer ^16, 34^. It is possible disrupting the scaffold would only have a modest effect, because the Vps5/Vps17 dimer remains intact and available to sort Vps10. Our data could be interpreted as providing indirect support for sorting of Vps10 by SNX-BAR proteins in yeast.

We note that interfaces observed in flat VPS35 dimers from mammalian chains look similar to the VPS35 dimer observed at the top of thermophilic yeast arches; the mammalian dimer is just flatter (Figure S9A). This places arch apexes closer to the membrane and causes the rest of the heterotrimer to adopt a different conformation, which leads to formation of flat chains instead of tall arches. Our multibody refinement revealed retromer exhibits inherent flexibility at multiple points, including a hinge point in the VPS35 solenoid; at VPS35/VPS35 dimer interfaces; and at VPS26/N-VPS35 interfaces. Taken together, these data support the idea that retromer is an adaptable and plastic scaffold that can adopt a range of conformations. Retromer may interact with SNX-BAR proteins by forming arches ^28^; SNX-BAR binding could promote or induce arch formation by organizing the scaffold. Modeling suggests flat chains could bind either SNX27 or SNX3, both of which lack BAR domains, in orientations placing lipid- or cargo-binding domains close to the membrane. Finally, although they are not well-resolved, the VPS26/VPS26 dimer observed in chain links (Figure S2) is very different from the yeast VPS26 dimer observed in the presence of Vps5^28^. Chains seem to form by tip-to-tip interactions between adjacent VPS26 subunits, as opposed to back-to-back interactions between N-VPS26 subdomains. Higher resolution structures for both binding modes will be required to understand these structures in more detail and to test assembly interfaces both *in vitro* and *in vivo*.

Why might retromer scaffold plasticity be advantageous for cells? Yeast retromer is an obligate pentamer, in which the heterotrimer assembles with Vps5/Vps17 dimers; both yeast sorting nexins contain BAR domains that recognize or drive membrane curvature. Mammalian retromer instead assembles with a variety of SNX proteins, including SNXs that lack BAR domains. Mammalian retromer also sorts cargoes to different destinations from a common origin. Thus, metazoan retromer may require plasticity to assemble distinct retromer complexes with specific sorting nexins to sort cargoes to different destinations.

An unresolved question is whether the tetramers we observe are relevant to retromer assembly. Tetramers are observed in micrographs (Figure S1) and in solution (Figure S3). Tetramers must interact via both VPS35 and VPS26-mediated interfaces. There are at least three possibilities that explain our observation of tetramers. First, they may be a biochemical artifact. We do not favor this interpretation, because we can specifically disrupt their formation with point mutations, but it formally remains a possibility. Second, tetramers may represent a cytosolic species that provides a retromer pool for rapid assembly on membranes in the presence of sorting nexins and cargo. Clathrin has been shown to assemble on membranes from pre-existing cytosolic pools^36, 37^, and we can conceptually consider retromer as a scaffold analogous to the clathrin cage. Finally, tetramers may represent another way to interact with membranes to sort cargoes; perhaps retromer could interact specific SNX proteins in this way. We cannot ascertain the orientation of VPS26 in our current tetramer reconstructions. We will require many more particles to generate higher resolution structures and ascertain tetramer architecture.

### Implications for human disease

The VPS35/VPS35 interfaces observed here and in the yeast cryoET structure have relevance to human disease. The VPS35 D620N mutation is linked to an autosomal dominant form of Parkinson’s disease ^38, 39^. Our structural data indicate D620 is located close to the acidic patch formed by E615, D616, and E617. This may suggest D620 plays an indirect role in maintaining the acidic surface required at the interface. Its mutation to asparagine may alter overall charge distribution and destabilize retromer assembly. This mutation has been proposed to interfere with the ability of VPS35 to associate with the WASH complex ^40, 41^. The WASH complex may favor interacting with a retromer dimer, and thus destabilizing the VPS35/VPS35 dimer interface would indirectly affect the ability of retromer to engage WASH. Alternatively, because this residue is located underneath the VPS35 dimer in the arch, it may play a role in engaging other binding partners.

## Materials and methods

### Reagents

Unless otherwise noted, all chemicals were purchased from Sigma (St. Louis, MO, USA).

### Molecular biology and cloning

Retromer constructs were generated in the labs of David Owen and Brett Collins and have been published previously ^20, 21^. We used a two-stage quick-change mutagenesis protocol adapted from Wang and Malcolm (Biotechniques 1999) to introduce point mutations into retromer plasmids to generate retromer electrostatic mutants. Briefly, mutagenic primers (Sigma) were created for the desired mutations. In the first step, two polymerase chain reactions (PCRs), with either the mutagenic 5’ or 3’ primer, were amplified around the plasmid. The two reactions were then combined in an additional PCR step, and the product was digested using Dpn I. Digested product was used to transform XL1 Blue (Agilent) competent cells, and colonies were sequenced using Sanger methods (Genewiz).

### Protein expression and purification

We expressed and purified recombinant retromer (wild-type and mutants) from *E. coli* as previously described ^21^. Briefly, retromer plasmids were transformed into BL21(DE3) Rosetta2 pLysS cells (Millipore). Cells were grown to OD_600_ between 0.8-1.0 and induced for 16-20 hours at 22°C with 0.4 mM IPTG. Cells were lysed by a disruptor (Constant Systems Limited). Protein was purified in 10 mM Tris-HCl (pH 8.0), 200 mM NaCl, 2 mM βME using glutathione sepharose (GE Healthcare). Protein was cleaved overnight using thrombin (Recothrom, The Medicines Company) at room temperature and batch eluted in buffer. Retromer was further purified by gel filtration on a Superdex S200 analytical column (GE Healthcare) into 10 mM Tris-HCl (pH 8.0), 200 mM NaCl, or dialysis into 20 mM HEPES pH 8.2, 50 mM NaCl, 2 mM DTT for cryoEM experiments.

### Negative stain grid preparation & screening

For screening of negatively stained retromer samples, 10 μl of retromer at concentrations between 5 and 10 μg/mL were applied to continuous carbon film on 400 square mesh copper EM grids (Electron Microscopy Sciences, Hatfield, PA) and washed twice with water. The grids were stained with 2% uranyl formate and 1% uranyl acetate and air dried overnight. The grids were screen on a ThermoFisher FEI Morgagni microscope operating at 100kV with a AMT 1k×1k CCD camera (Vanderbilt University CryoEM Facility) to verify protein quality. The grids were imaged on a ThermoFisher FEI Tecnai F20 operating at 200 kV with a 4kx4k CCD camera (Vanderbilt University CryoEM Facility).

### Cryo-EM grid preparation and data collection

For cryo-electron microscopy, retromer at a concentration of 100 µg/ml was applied to freshly glow discharged CF-2/2-2C C-Flat grids (Protochips, Morrisville, NC), and the grids were vitrified in liquid ethane using either a ThermoFisher FEI MarkIII or MarkIV Vitrobot. Data were collected from 2779 micrographs at the National Resource for Automated Molecular Microscopy (NRAMM) in two different data collection sessions using ThermoFisher FEI Titan Krios microscopes operating at 300keV. The first data collection used a Titan Krios equipped with a spherical aberration (Cs) corrector; a Volta phase plate; a Gatan BioQuantum energy filter; and a Gatan K2 Summit camera. The second data collection used a similar Krios instrument but without a Cs corrector. The nominal magnification used during each data collection was 105,000x and 130,000x, respectively. The effective pixel size was 1.096Å/pix for the first data collection and 1.06Å/pix for the second data collection. The total electron dose during the two data collection sessions was between 69 and 74 e^-^/A^2^. Table S1 contains a data collection summary.

### Single particle cryo-EM image & data processing

All images were motion corrected using MotionCor2^42^. Because we used different Krios microscopes in two data collection sessions, micrographs from the second data collection were rescaled to match the 1.096Å/pix pixel size from the first data collection using an NRAMM script written for MotionCor2. The CTF of each micrograph was determined using Gctf ^43^; defocus values for the data varied between −0.7 and −4.4μm. RELION-2^44^ and RELION-3^45^ were used for image processing.

For each dataset, we manually selected several thousand particles and performed an initial 2D classification to produce templates for autopicking. Autopicking for the heterotrimer, dimer, flat chain, tetramer, and curved and flat substructures identified 249,562 particles from the first Krios dataset and 190,084 particles from the second dataset. Autopicking for the chainlink and centered link chain identified 301,950 particles from the first Krios dataset and 141,167 particles from the second dataset. Multiple rounds of 2D classification were performed to separate the different species into distinct particle sets. Initial models for 3D classification were generated from models generated by earlier experiments, filtered to 60 Å resolution. Detailed methods are provided in Extended EM data processing methods, and statistics are provided in Table S2.

### Model building & docking

An initial model of mammalian retromer heterotrimer was produced by combining partial X-ray crystallographic structures of retromer subunits (PDB: 2R17, 5F0J). Models for VPS29 and the VPS35 C-terminus were obtained from PDB 2R17, while models for N-VPS35 and VPS26A were obtained from PDB 5F0J. We omitted a flexible unstructured loop (amino acids 470-482) that is absent in all crystal structures. Rigid-body docking and map visualization were performed using Chimera (Pettersen *et al.*, 2004) using Chimera Fit in Map routine.

### SEC MALS

Size-exclusion chromatography coupled to multi-angle laser light scattering (SEC-MALLS) was performed on an ÄKTA Purifier FPLC system (GE Healthcare) in-line with an Agilent Technologies 1200 Series refractive index Detector and a Dawn Helios 8+ MALS unit (Wyatt). 100 µL of sample at 15 µM was injected from a static loop beginning at the 0 ml point of each run onto a Superdex 200 increase 10/300 GL column (GE Healthcare) at a flow rate of 0.5 ml/min. Samples were assayed at 25°C in 20 mM Tris-HCl (pH 8.5), 2 mM DTT, and 8 mM NaN_3_ at two different salt concentrations (50 mM NaCl and 500 mM NaCl). Refractive index and light scattering data were recorded and analyzed using ASTRA 6.1 (Wyatt) software. Molecular masses were calculated across distinct eluted peaks using the Zimm algorithm with a dn/dc value of 0.185 ml/g.

### Dynamic light scattering

Dynamic light scattering experiments were performed using a Wyatt Technology DynaPro NanoStar instrument. Retromer samples were assayed in 20 mM Tris-HCl (pH 8.5), 2 mM DTT at two different salt concentrations (50 mM NaCl and 500 mM NaCl). 10 µL of sample at 7 µM was equilibrated at 25°C for 5 minutes prior to data acquisition in a disposable 4 µL cyclic olefin copolymer cuvette. Twenty acquisitions at 5s intervals were recorded for each sample and fitted with the coils shape model using the Wyatt Dynamics software.

### Yeast strains

Standard media and techniques for growing and transforming yeast were used. Yeast genes knockout was performed using a PCR toolbox ^46^. VPS26VPS35 double knockout strains were generated from a *ΔVPS35* strain ^33^. The double *ΔVPS35/ ΔVPS26* knockout strain was confirmed by PCR-based genotyping. Details on yeast strains are provided in Table S3.

### Carboxypeptidase Y secretion assay

Retromer mutant yeast cells were 10-fold serial diluted and spotted on YPD plates at 26°C. The cells were grown for 24 hr prior to being overlaid with a nitrocellulose membrane. The operation is carefully performed to prevent air bubble from being trapped under the membrane. The plates were grown for an additional day. The nitrocellulose membranes were washed several times with a gentle flow of deionized water and then placed in PBS-T buffer (standard phosphate-buffered saline with 0.1% (v/v) Tween-20) until ready to use. The membrane was blocked in Odyssey Blocking Buffer (LI-COR Biosciences) for 1 hr at room temperature then incubated with anti-CPY antibodies (Invitrogen) at1:1000 in blocking buffer overnight. After washing with PBS-T, the membrane is incubated with HRP-goat anti-mouse IgG antibody (Invitrogen) (1:10,000 in PBS-T +5% non-fat milk) for 1 hr at room temperature. The membrane was washed by PBS-T and then developed by Amersham ECL Western Blotting Detection Reagent (GE Healthcare) and imaged with AI600 Chemiluminescent Imager system (GE Life Sciences). The dots intensity was quantified by ImageStudio (GE Life Sciences). Statistical differences were determined using a one-way ANOVA on the means of at least three independent experiments using GraphPad Prism (GraphPad Software). Probability values of less than 0.05, 0.01 and 0.001 were used to show statistically significant differences and are represented with *, ** or *** respectively.

### Data availability

EM electron density maps were deposited in the EMDB with accession numbers EMD-VV, EMD-WW, EMD-XX, EMD-YY, and EMD-ZZ corresponding to the retromer heterotrimer, dimers, chains, and tetramers. Sub-structure maps for the flat and curved VPS35 dimers were deposited as EMD-XXX and EMD-YYY. Pseudo-atomic coordinates for the heterotrimer and flat VPS35/VPS35 sub-structures were deposited with PDB accession codes XXXX and YYYY.

## Acknowledgements

We sincerely thank Bridget Carragher and Clint Potter for their generosity in providing access to excellent instrumentation. Electron microscopy data collection was performed at the Simons Electron Microscopy Center and National Resource for Automated Molecular Microscopy located at the New York Structural Biology Center, supported by grants from the Simons Foundation (349247) and the NIH National Institute of General Medical Sciences (GM103310). We thank Michael Sheedlo, Rick Baker, and Mike Cianfrocco for helpful feedback and discussions on data processing; and Brett Collins and David Owen for helpful discussions and sharing unpublished data. AK, BX, RB, MF and LPJ are supported by NIH R35GM119525. LPJ is a Pew Scholar in the Biomedical Sciences, supported by the Pew Charitable Trusts. Initial screening for EM data collection was undertaken on a ThermoFisher FEI Polara and TF20 microscopes at the Center for Structural Biology CryoEM facility (V-CEM) at Vanderbilt University. We thank Dr. Scott Collier for his support at the facility.

## Author contributions

AKK, BX, RB, JW, MNF, and LPJ purified proteins, generated mutants, and undertook *in vitro* experiments. EB, HW, and TN assisted with data collection, interpretation, and processing. BX, PX, TG, and LPJ designed and conducted yeast experiments. AKK and LPJ processed data, generated models, and wrote the manuscript with input and feedback from all authors. LPJ conceived and designed the project.

---

**Figure S1.**
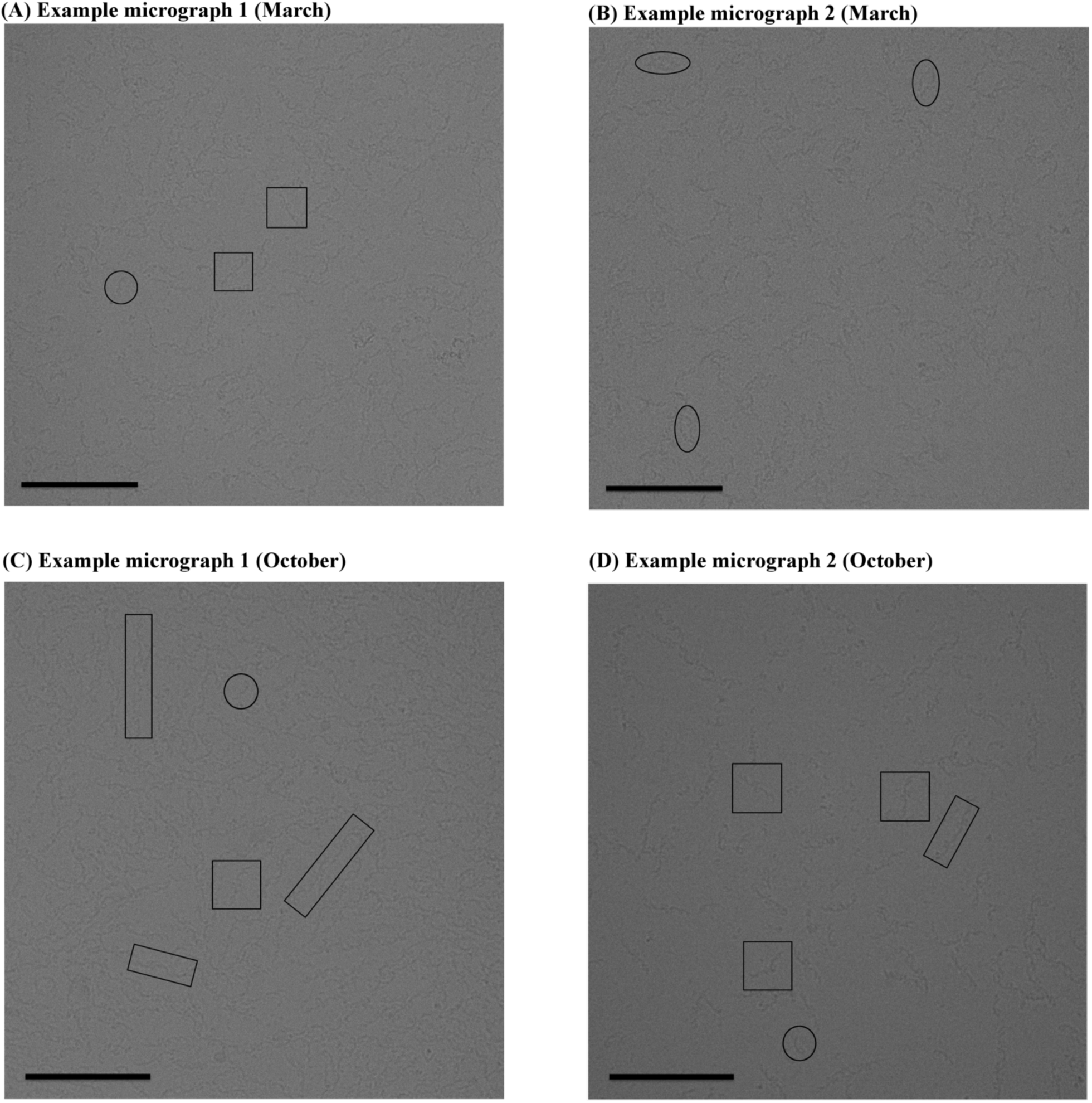
Retromer samples for cryo-electron microscopy. Representative micrographs of retromer in vitrified ice. (A) and (B) show two micrographs from data collection #1 (March 2018). (C) and (D) show two micrographs from data collection #2 (October 2018). Heterotrimers are marked in circles; dimers are marked in squares; chains are marked in rectangles; and tetramers are marked in ovals. Scale bars represent 100 nm.

**Figure S2.**
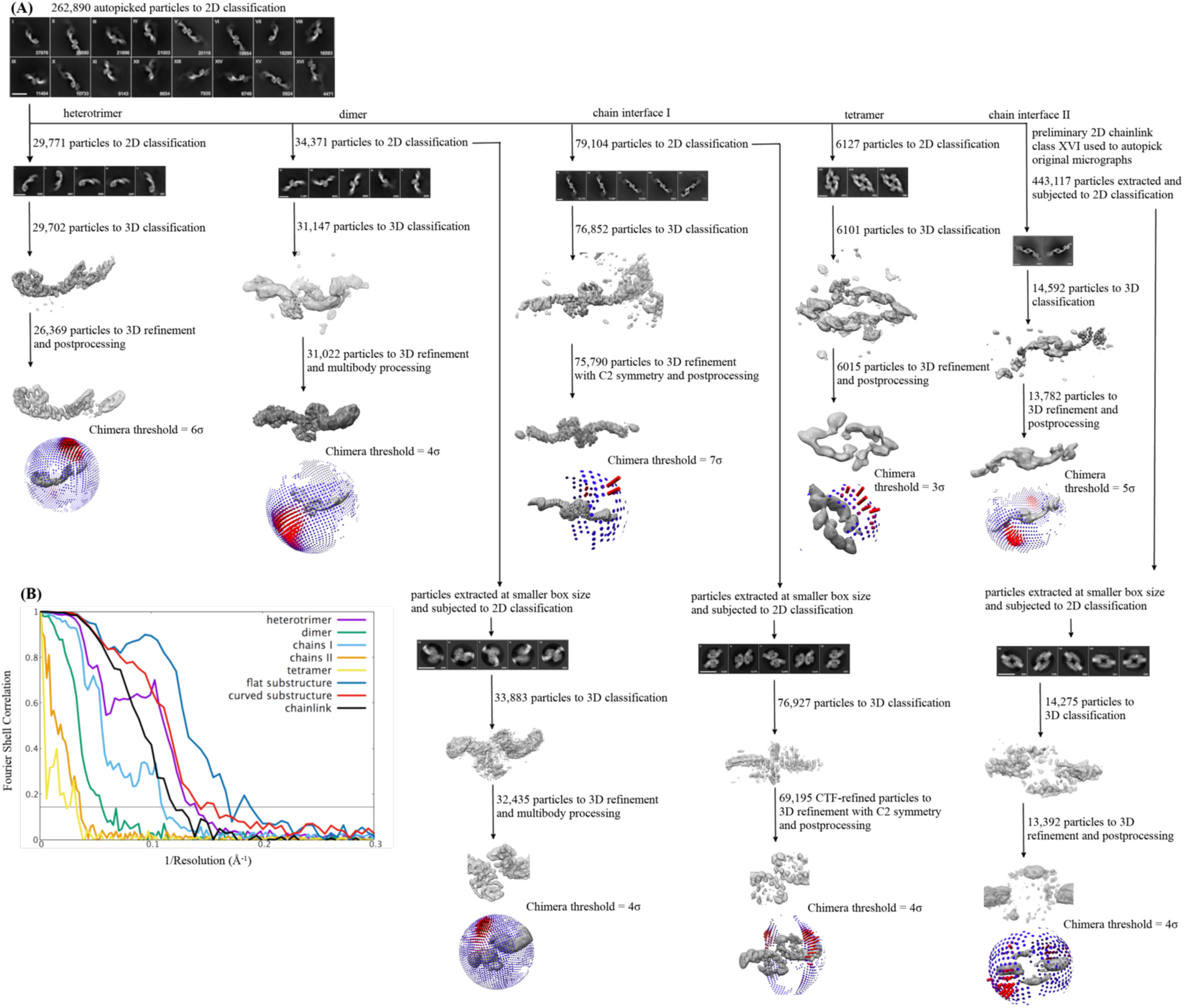
Image and data processing work flow. (A) The cryo-EM analysis workflow is shown. Particles were initially auto-picked from two combined datasets and then separated into 2D classes based on biochemical species (heterotrimer, dimers, chains, and tetramers). 3D reconstructions were generated for each species. Flat and curved VPS35/VPS35 sub-structures were generated from chains and dimers, respectively, by masking VPS26 and N-VPS35. (B) Fourier Shell Correlation (FSC) plots for each structure and sub-structure. The grey dotted line marks the “gold standard” 0.143 cut-off, which we use for reported resolution values in the manuscript.

**Figure S3.**
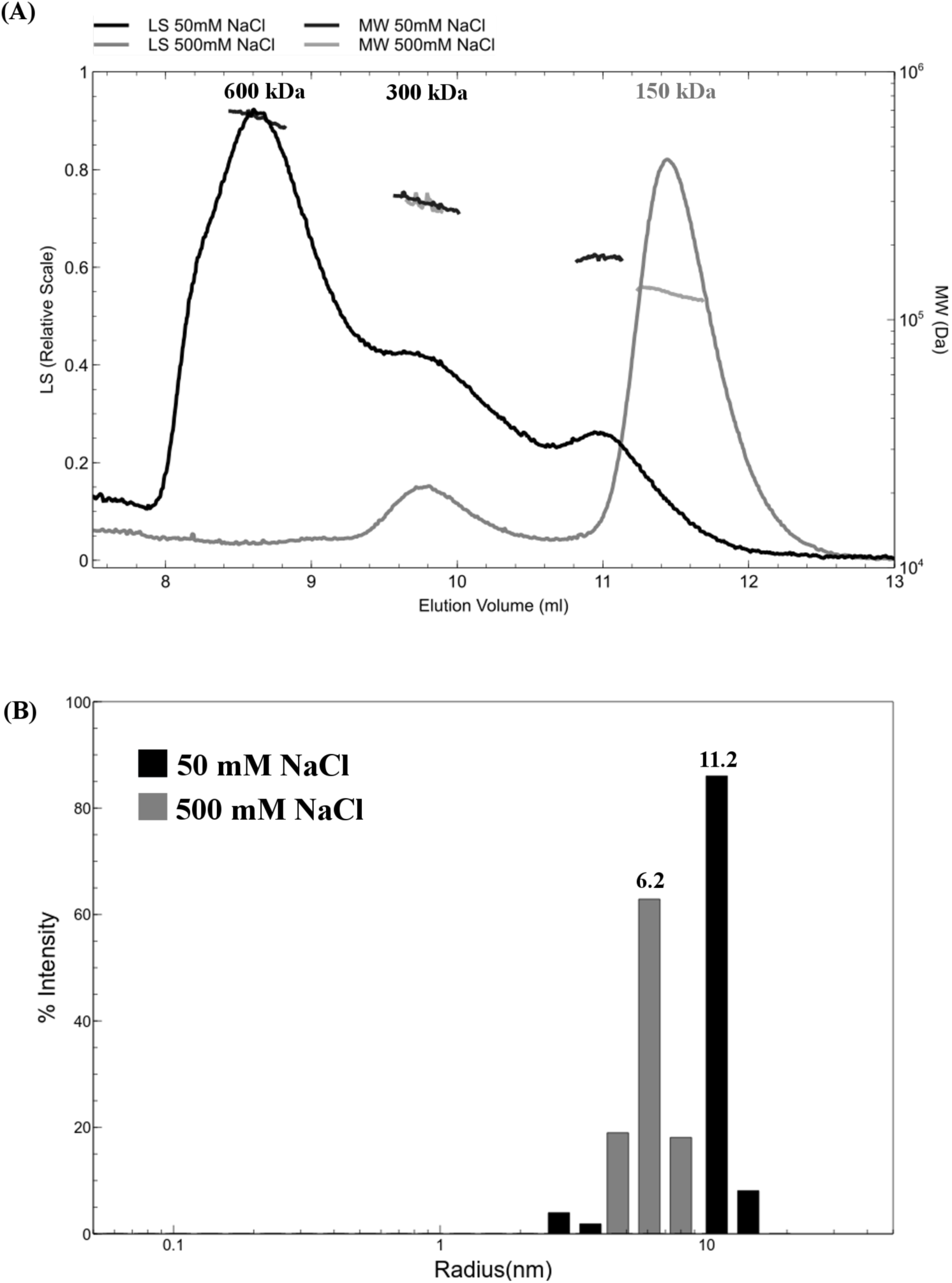
Mammalian retromer forms oligomers in solution in a salt-dependent manner. (A) At 500 mM NaCl (grey trace), the main retromer peak elutes from a size exclusion column at a volume consistent with one copy of the VPS26/VPS35/VPS29 heterotrimer (~150 kDa). A small population of retromer elutes in a second peak as a “dimer of trimers” (~300 kDa). At 50 mM NaCl (black trace), the peak profile shifts: the predominant peak is now consistent with four copies of the heterotrimer (600 kDa); a second peak or shoulder is consistent with two copies, and the third is consistent with one copy. (B) Dynamic light scattering reveals that retromer particles in low salt (50 mM NaCl, grey bars) have approximately double the average radius compared to retromer in high salt (500 mM NaCl, black bars). This is consistent with SEC MALS data in (A) indicating retromer forms oligomers in solution.

**Figure S4.**
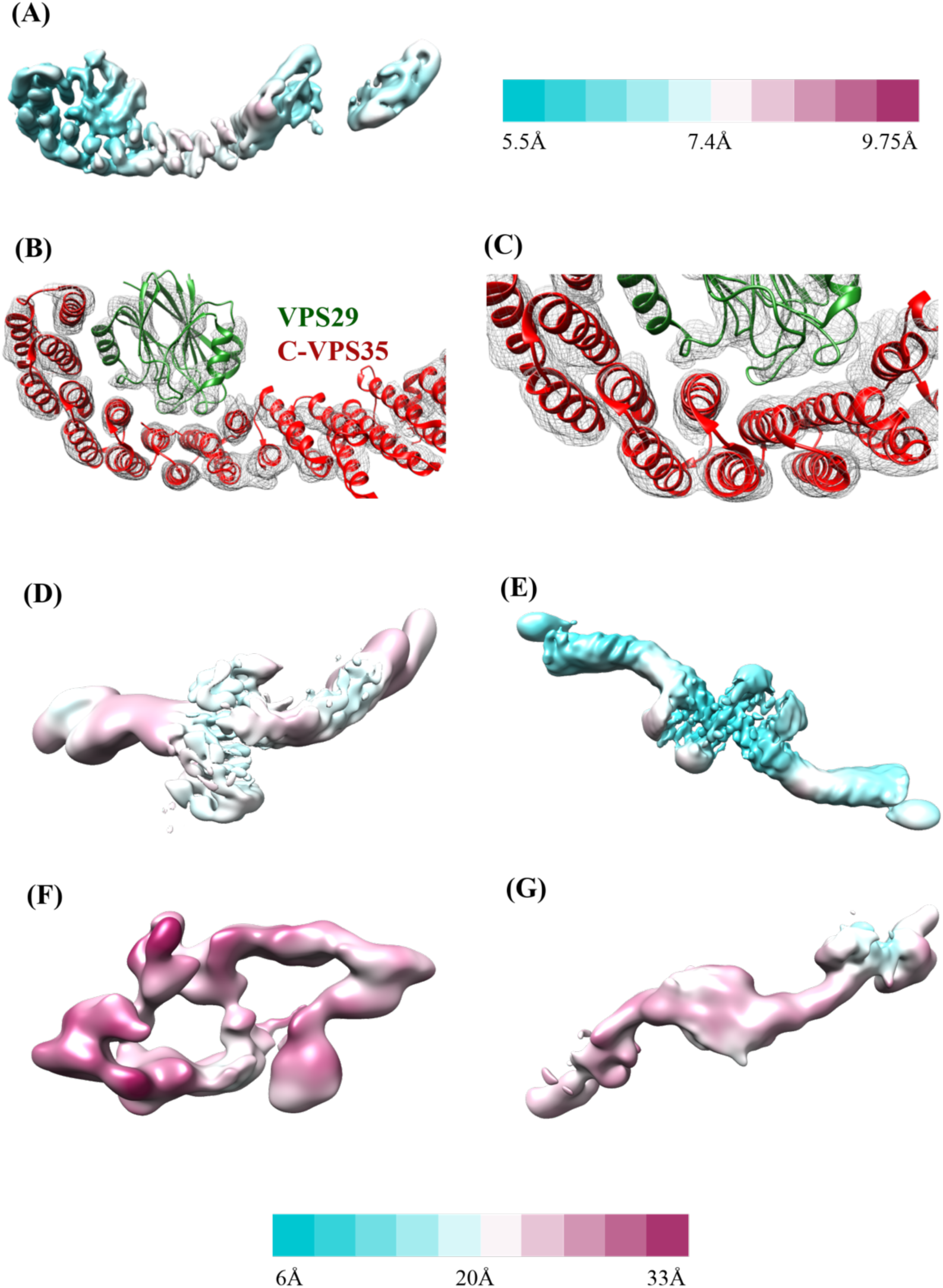
Local resolution and features. (A) Retromer heterotrimer resolution range across the model with legend shown to right (5.5-9.8 Å). (B), (C) Two different views of heterotrimer map quality to show model fitting into α-helices (contoured at 6σ). (D-G) Resolution range for dimer (D); VPS35-centered chain interface I (E); tetramer (F); and VPS26-centered chain interface II (G). Legend shown below from 6 (cyan) to 33 (magenta) Å

**Figure S5.**
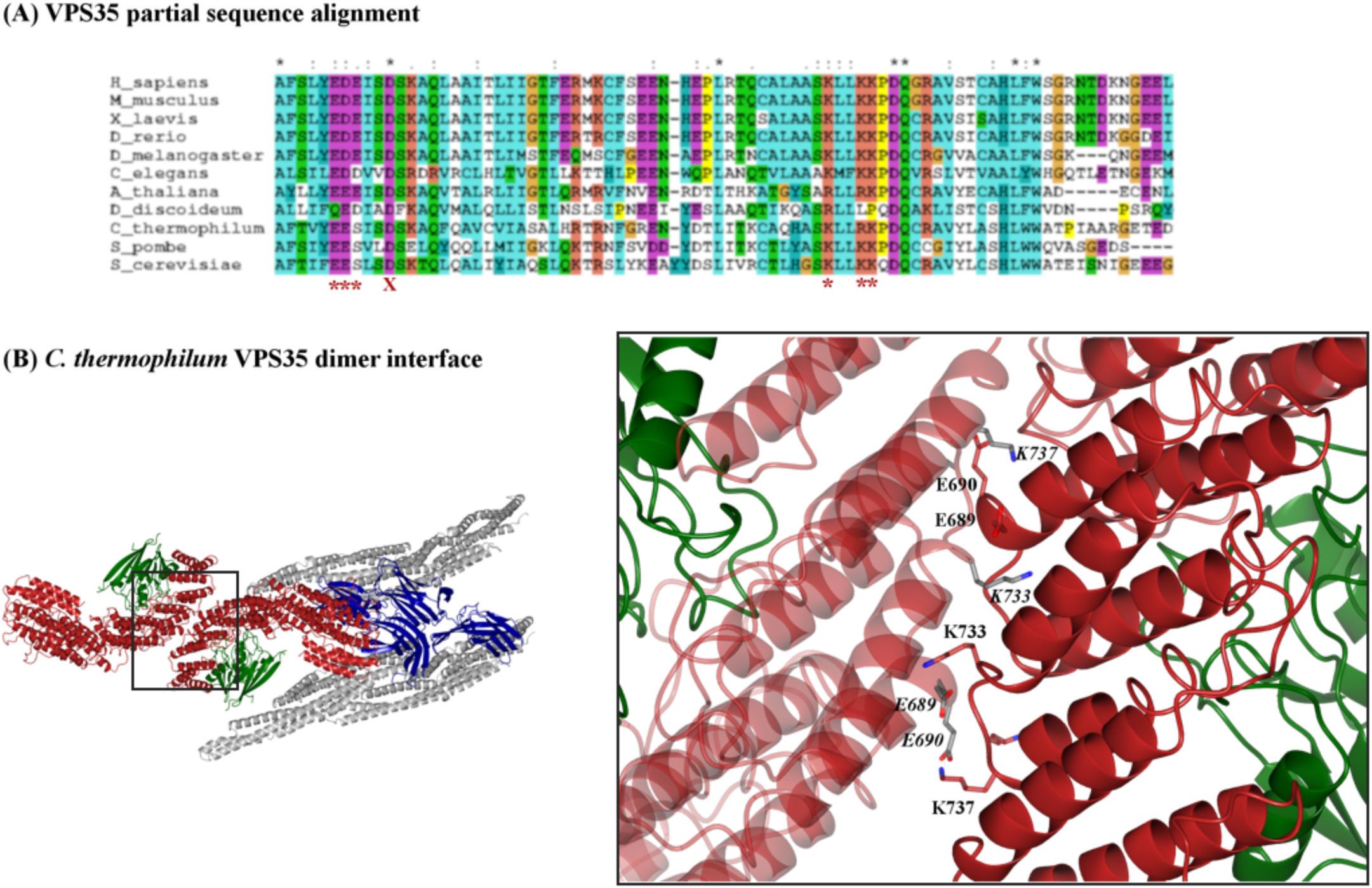
Conservation in the VPS35 C-terminus. (A) Sequence alignments were generated in Clustal Omega, and a partial alignment corresponding to *M. musculus* amino acid residues 610-690 are shown here for clarity. The following species were used to represent diversity across eukaryotes in the full-length alignment: *S. cerevisiae, S. pombe, C. thermophilum, D. discoideum, A. thaliana, C. elegans, D. melanogaster, D. rerio, X. laevis, M. musculus*, and *H. sapiens.* Red asterisks below the alignment mark conserved residues; these residues were mutated to generate the electrostatic mutant described in the main text. The red X marks residue D620, a residue implicated in late-onset Parkinson’s when mutated to asparagine. (B) Conserved acidic and basic residues are found in the VPS35/VPS35 dimer interface of the *C. thermophilum* cryo-ET structure. The left-hand view shows a top-down view of Vps5/retromer arches. The VPS35 dimer interface shown in the boxed inset. E689, E690, K733, and K737 are the thermophilic yeast equivalents of mouse residues E615, D616/E617, K659, and K662.

**Figure S6.**
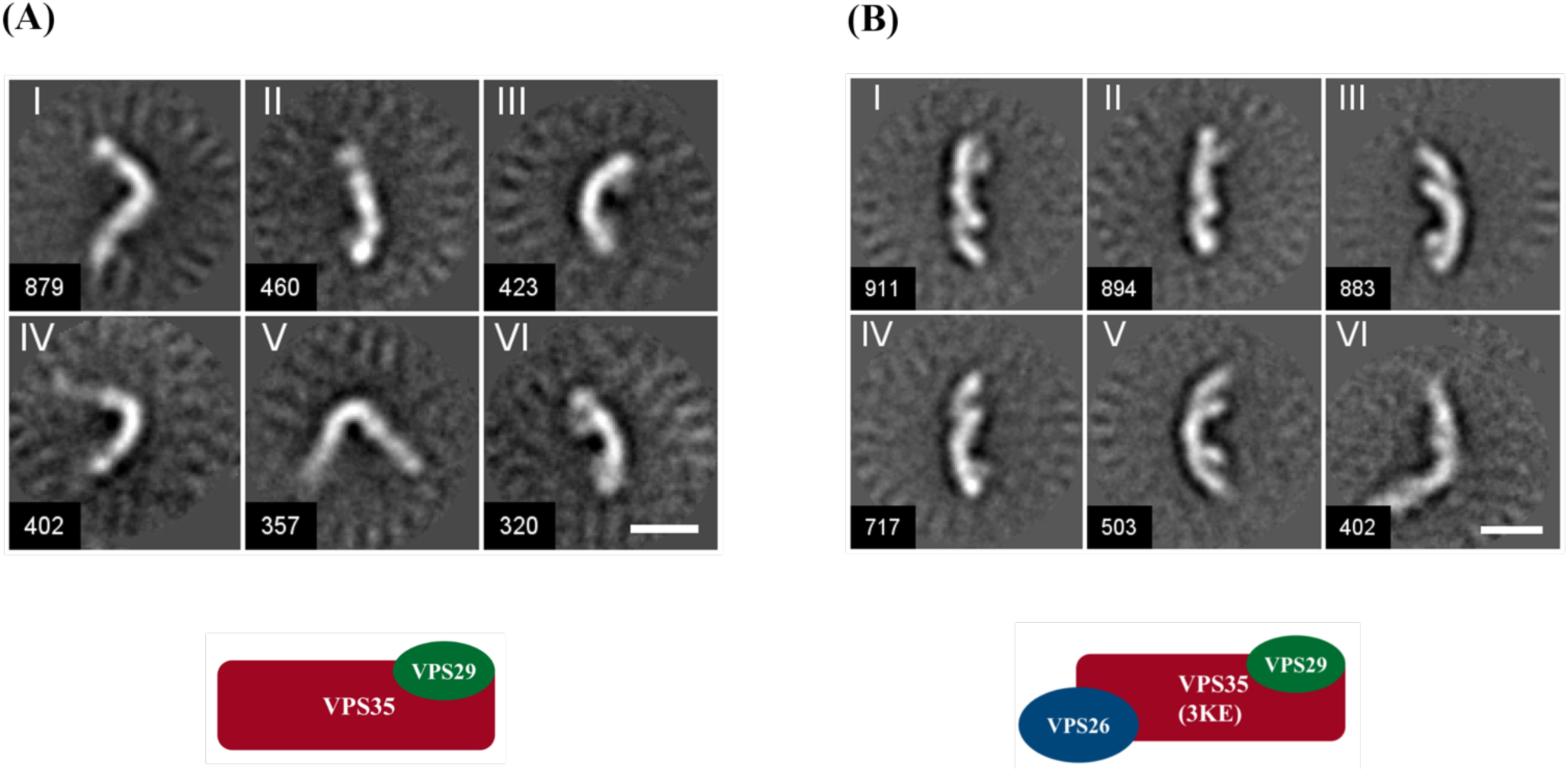
Retromer mutants exhibit assembly defects *in vitro*. (A) Representative 2D class averages of negatively stained VPS35/VPS29 sub-complex particles. The VPS35/VPS29 sub-complex forms dimers but exhibits greater flexibility when VPS26 is absent. (B) Representative 2D class averages of negatively stained partial retromer electrostatic mutant, 3KE (K659E/K662E/K663E). Scale bars represent 10 nm.

**Figure S7.**
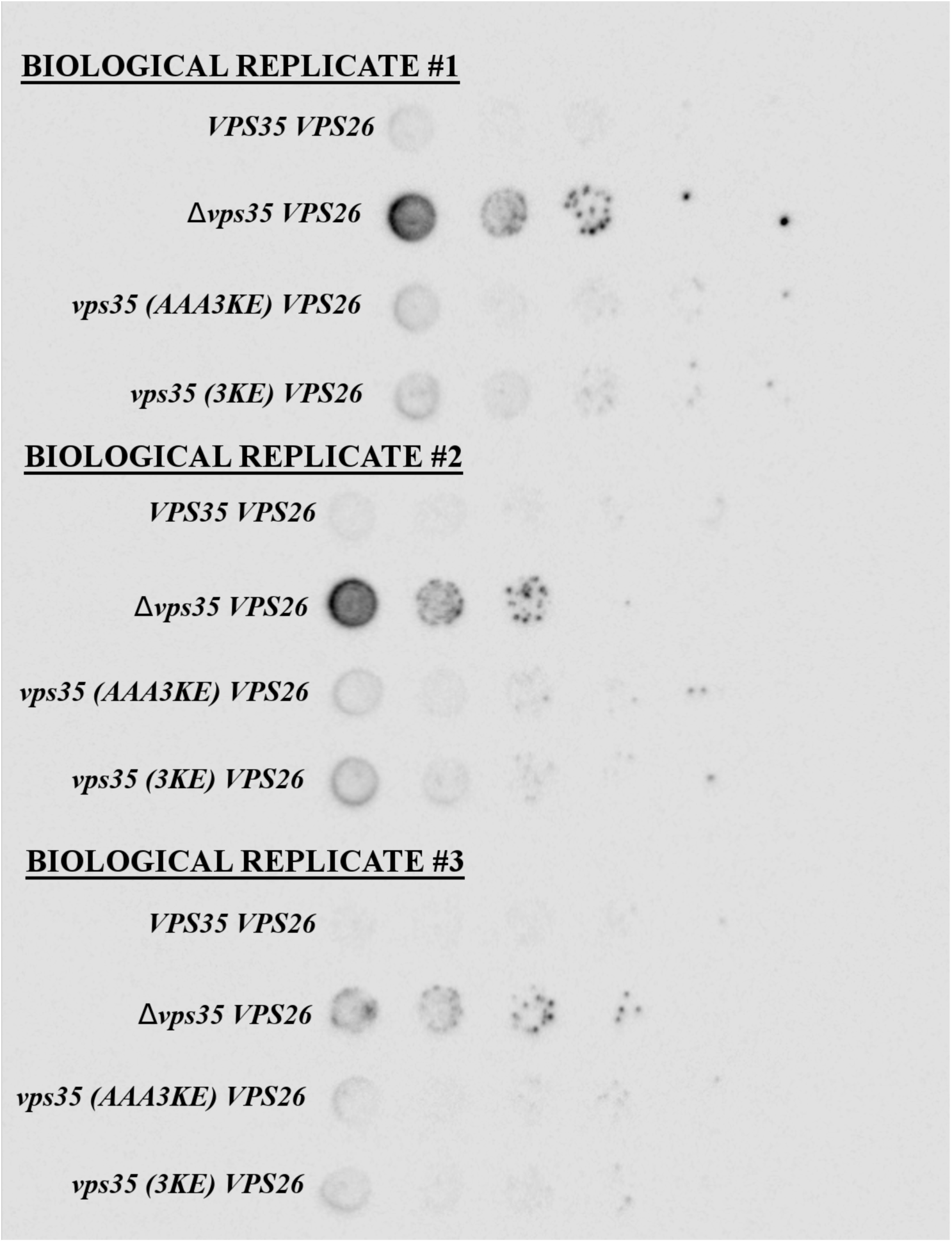
Carboxypeptidase Y (CPY) sorting assay in budding yeast. Three biological replicates are shown from the CPY secretion assay. The box from replicate #1 represents the cropped blot shown in main Figure 4.

**Figure S8.**
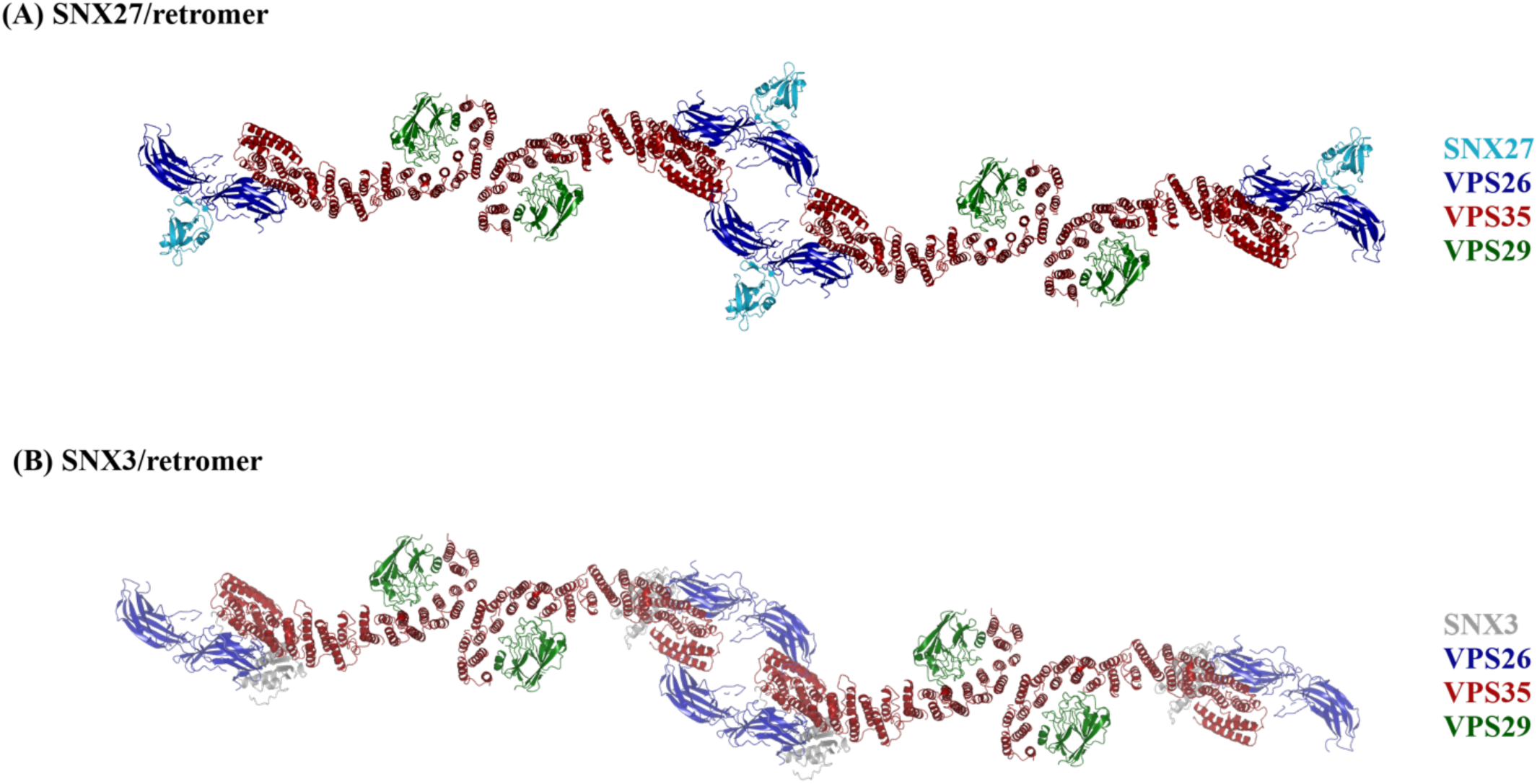
Models of flat retromer chains with mammalian sorting nexins. (A) Top down view of SNX27/retromer chains with SNX27 PDZ domain shown in cyan (PX and FERM domains are omitted). The SNX27 PDZ domain points out and down from the VPS26 arrestin saddle in this model. (B) Top down view of SNX3/retromer chains, with SNX3 PX domain shown in grey. The SNX3 PX domain points down towards a membrane in this model. VPS26 is shown in blue, VPS35 in red, and VPS29 in green.

**Figure S9.**
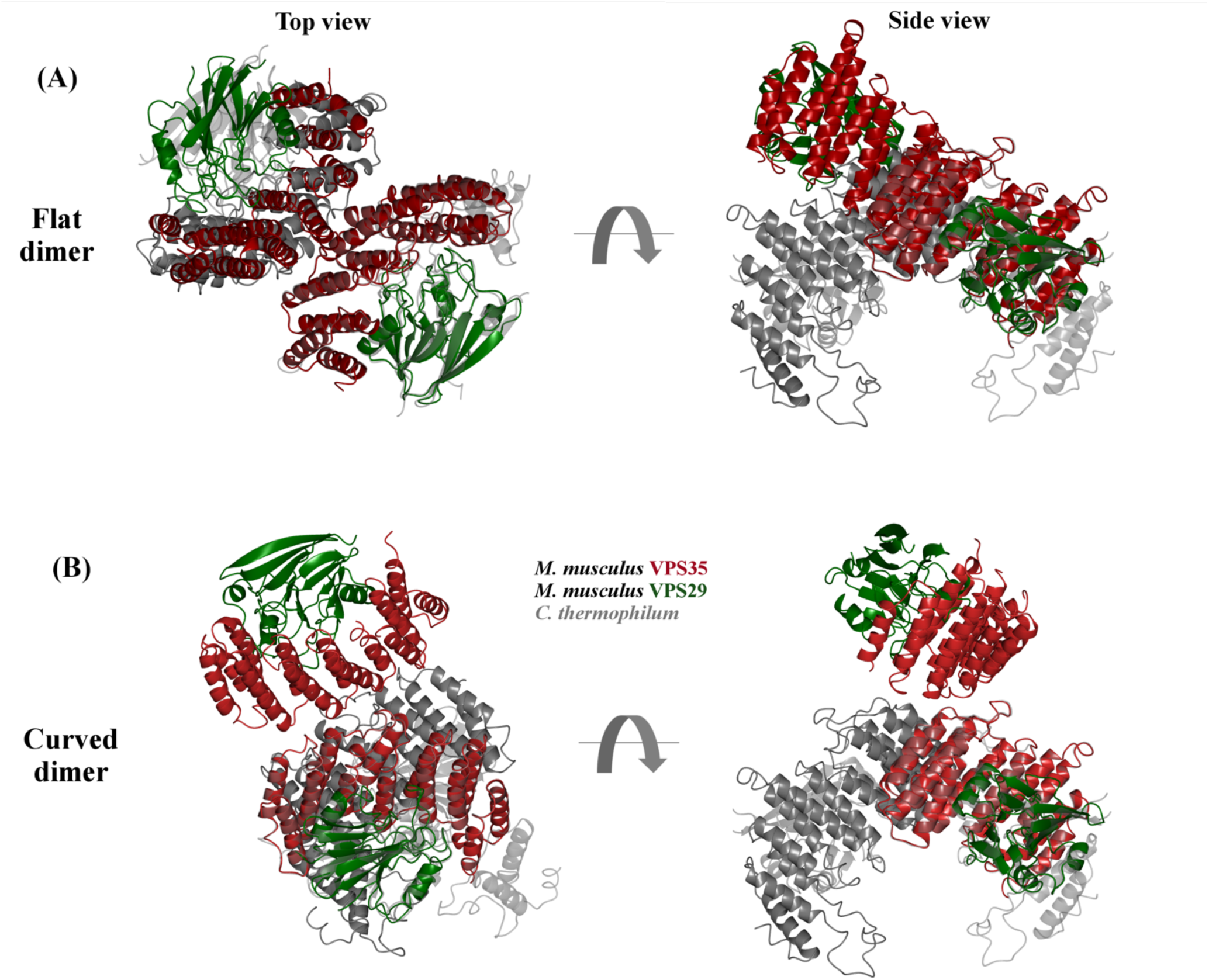
Comparison of mammalian and thermophilic yeast dimers. The flat (A) and curved (B) mammalian VPS35 dimers (VPS35 in red, VPS29 in green) were superposed onto the thermophilic yeast VPS35 dimer (grey) that forms the top of retromer arches when reconstituted on membranes with Vps5. Two views rotated by 90° are shown. Top views (left-hand column) are shown looking down onto the membrane from above. Side views (right-hand column) represents the apex of an arch in the thermophilic yeast structure.

**Table S1.**
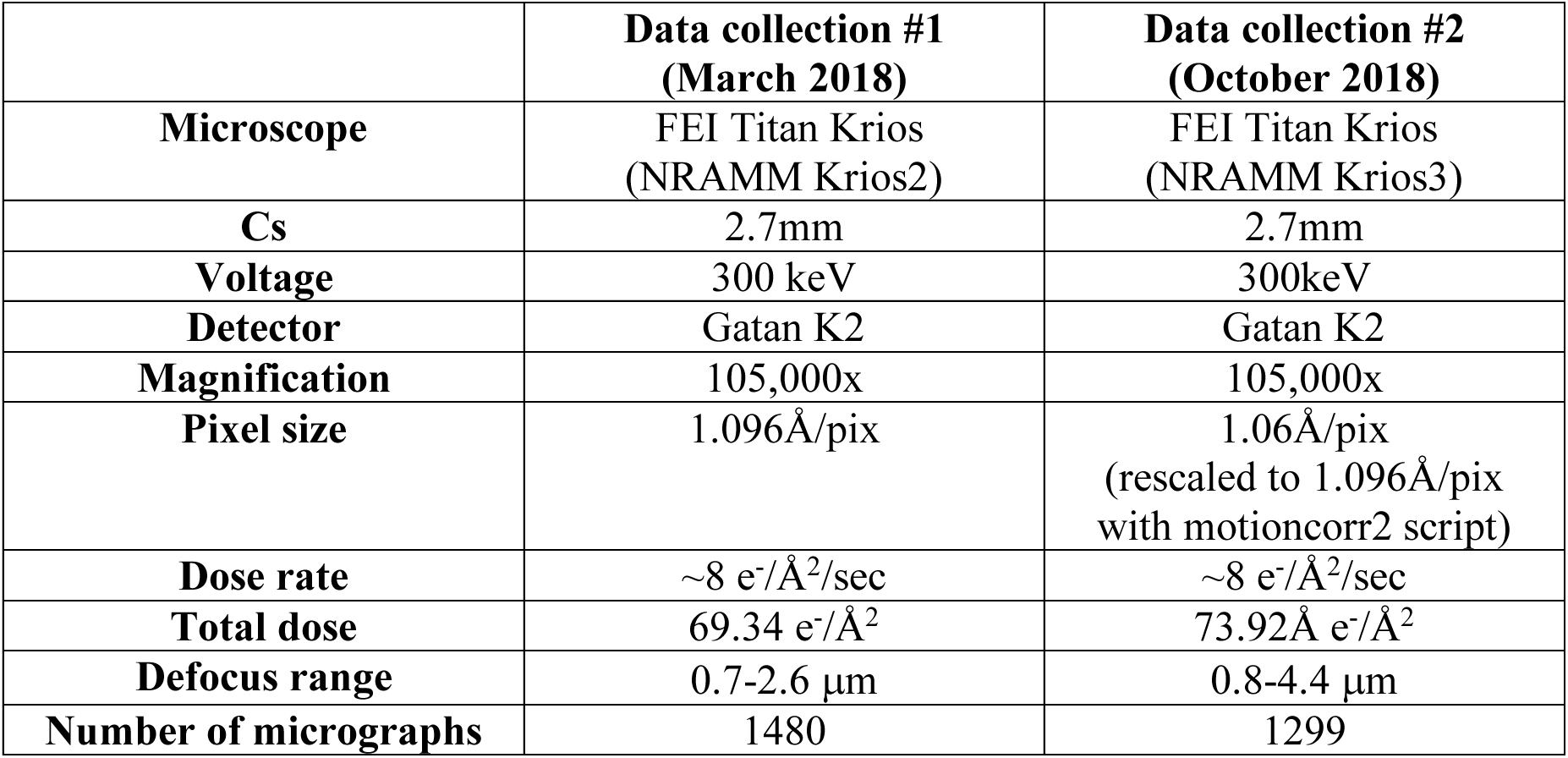
Data collection parameters.

|  | **Data collection #1**<br>**(March 2018)** | **Data collection #2**<br>**(October 2018)** |
| --- | --- | --- |
| **Microscope** | FEI Titan Krios<br>(NRAMM Krios2) | FEI Titan Krios<br>(NRAMM Krios3) |
| **Cs** | 2.7mm | 2.7mm |
| **Voltage** | 300 keV | 300keV |
| **Detector** | Gatan K2 | Gatan K2 |
| **Magnification** | 105,000x | 105,000x |
| **Pixel size** | 1.096Å/pix | 1.06Å/pix<br>(rescaled to 1.096Å/pix<br>with motioncorr2 script) |
| **Dose rate** | ~8 e<sup>-</sup>/Å<sup>2</sup>/sec | ~8 e<sup>-</sup>/Å<sup>2</sup>/sec |
| **Total dose** | 69.34 e<sup>-</sup>/Å<sup>2</sup> | 73.92Å e<sup>-</sup>/Å<sup>2</sup> |
| **Defocus range** | 0.7-2.6 μm | 0.8-4.4 μm |
| **Number of micrographs** | 1480 | 1299 |

**Table S2.** Data processing and refinement statistics. Data processing statistics are reported for all structures and sub-structures. The heterotrimer and flat VPS35/VPS35 sub-structure underwent real space refinement in PHENIX.

|  | heterotrimer | dimer | curved<br>VPS35/ VPS35<br>sub-structure | chain<br>interface<br>I | flat<br>VPS35/VPS35<br>sub-structure | tetramer | chain<br>link | chain<br>interface<br>II |
| --- | --- | --- | --- | --- | --- | --- | --- | --- |
| Total particles (autopicked) | 439,646 | 439,646 | 439,646 | 439,646 | 439,646 | 439,646 | 443,117 | 443,117 |
| Box size (Å) | 252x252 | 395x395 | 180x180 | 395x395 | 180x180 | 329x329 | 180x180 | 482x482 |
| Particles in 2D classification | 29,771 | 31,202 | 34,156 | 76,927 | 78,994 | 6126 | 15,522 | 15,234 |
| Particles in final 3D model | 26,369 | 31,022 | 32,435 | 75,790 | 69,195 | 6015 | 13,392 | 13,782 |
| Symmetry | C1 | C1 | C1 | C2 | C2 | C1 | C1 | C1 |
| Map resolution<br>(FSC 0.143 in RELION) | 6.0 Å | 14.6 Å | 5.5 Å | 7.1 Å | 5.0 Å | 27.4 Å | 6.2 Å | 18.5 Å |
| B-factor | -212.361 | -- | -57.878 | -188.756 | -55.7066 | -- | -298.916 | -212.361 |
| EMDB accession code | XXXX | XXXX | XXXX | XXXX | XXXX | XXXX | XXXX | XXXX |
| ***Refinement*** |  |  |  |  |  |  |  |  |
| Program | PHENIX |  |  |  | PHENIX |  |  |  |
| Number atoms | 9793 |  |  |  | 7563 |  |  |  |
| Molprobity score | 2.11 |  |  |  | 2.25 |  |  |  |
| Molprobity clash score | 17.05 |  |  |  | 20.8 |  |  |  |
| *RMSD from ideal* |  |  |  |  |  |  |  |  |
| Bond length (Å) | 0.006 |  |  |  | 0.006 |  |  |  |
| Bond angles (°) | 1.067 |  |  |  | 1.067 |  |  |  |
| *Ramachandran plot* |  |  |  |  |  |  |  |  |
| Favored (%) | 94.5 |  |  |  | 93.2 |  |  |  |
| Allowed (%) | 5.5 |  |  |  | 6.8 |  |  |  |
| Outliers (%) | 0.0 |  |  |  | 0.0 |  |  |  |
| Rotamer outliers (%) | 0.55 |  |  |  | 1.3 |  |  |  |
| CC model-vs-map (masked) | 0.70 |  |  |  | 0.64 |  |  |  |
| Map resolution (FSC 0.143) | 5.8 Å |  |  |  | 5.3 Å |  |  |  |
| PDB ID | YYYY |  |  |  | YYYY |  |  |  |

**Table S3.**
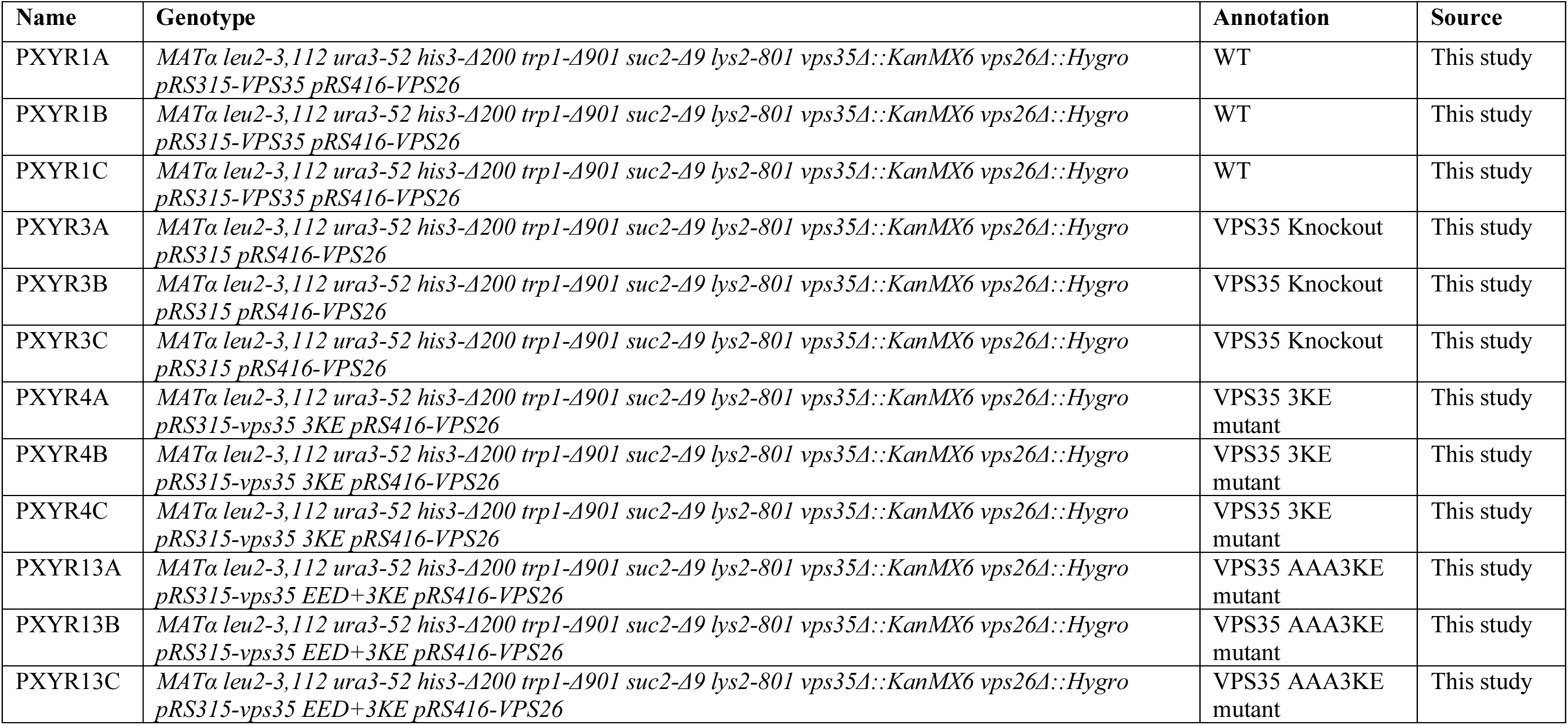
Yeast strains used in this study.

## Extended EM data processing methods

RELION-2^1^ and RELION-3^2^ were used for all EM image processing unless otherwise indicated.

2D classification of several thousand manually picked particles from our two datasets yielded templates for autopicking. Autopicking of the first Krios dataset identified 249,562 particles; autopicking of the second dataset identified 190,085 particles. These 439,646 particles were subjected to initial 2D classification, and 262,890 particles were retained. These particles were divided into five different streams for image processing: heterotrimer; dimer and curved sub-structure; chain interface I and flat substructure; chain interface II and chain link substructure; and tetramers. Numerous rounds of 2D classification were required to separate particles that appeared similar in 2D (for example, the dimer and tetramer particles).

### Heterotrimer

2D classification of the heterotrimer particle began with 29,771 particles at a box size of 252×252 Å. Multiple rounds of 2D classification and re-centering of these particles yielded 29,702 particles suitable to continue to 3D classification. A preliminary 3D model generated using only particles from the first dataset was filtered to 60 Å and used as an input model for 3D classification; 6 output classes were used. 26,369 particles from 3D classification were selected to continue to 3D refinement and postprocessing. The final postprocessed heterotrimer model had a resolution of 6.0 Å and a B-factor of −212.361. Multibody processing of the heterotrimer was performed using three different masks: one for VPS26; one for the N-terminus of VPS35 (representing the first 15 α-helices); and the C-terminus of VPS35 plus VPS29.

### Dimer

2D classification of the dimer particle began with 34,371 particles at a box size of 395×395 Å. Numerous rounds of 2D classification were required to separate the dimer structure from the flat chain (chain interface I), and 31,147 dimer particles were selected to continue to 3D classification. A preliminary 3D model generated using only particles from the first dataset was filtered to 60 Å and used as an input model for 3D classification; 6 output classes were used. 31,022 particles from 3D classification were selected to continue to 3D refinement. The final dimer model had a resolution of 14.6 Å, and this model did not benefit from postprocessing. Instead, multibody refinement was used to characterize the flexibility observed in these particles. Multibody refinement of the dimer using six independent masks (the same masks as described for heterotrimer particle, with one set of masks for each leg of the dimer) produced six different partial structures with resolutions ranging from 9.0-17.0 Å.

### Curved VPS35/VPS35 substructure

Processing of the curved sub-structure began with re-extraction of 34,371 dimer particles at a box size of 180×180 Å. Repeated rounds of 2D classification with a mask of 149 Å were performed. 33,883 particles were selected to continue to 3D classification, and a preliminary 3D model generated using only particles from the first dataset was filtered to 60 Å and used as input; 6 output classes were used. 32,435 particles from 3D classification were selected to continue to 3D refinement and postprocessing. The final postprocessed curved substructure model had a resolution of 5.5Å and a B-factor of −57.878. Multibody refinement of the curved substructure used two masks, one for each copy C-VPS35/VPS29 in the sub-structure. Postprocessing of these separate partial structures produced one copy at a resolution ~4.0 Å; the second copy was less well resolved at 6.0 Å.

### Chain interface I

For computational reasons, chain particles were separated into two units (called chain interface I and chain interface II) for data processing; the box size for a longer chain encompassing both interfaces is impractical. 2D classification of the chain interface I particle (centered on the VPS35/VPS35 interface) began with 79,104 particles at a box size of 395×395 Å. Numerous rounds of 2D classification were required to separate the chain interface I structure from the dimer. 76,852 particles were selected to continue to 3D classification using C2 symmetry, and a preliminary 3D model generated using only particles from the first dataset was filtered to 60 Å and used as input; 6 output classes were used. 75,790 particles were selected for 3D refinement and postprocessing. The final postprocessed model had a resolution of 7.1 Å and a B-factor of - 188.756.

### Flat substructure

Processing of the flat substructure began with re-extraction of 79,104 chain interface I particles at a box size of 180×180Å. Repeated rounds of 2D classification with a mask of 149 Å were performed. 76,927 particles were selected to continue to 3D classification using C2 symmetry, and a preliminary 3D model generated using only particles from the first dataset was filtered to 60 Å and used as input; 6 output classes were used. 69,195 particles selected from 3D classification were subjected to CTF refinement, then continued to 3D refinement and postprocessing. The final postprocessed flat substructure model had a resolution of 5.0Å and a B-factor of −55.7066.

### Chain interface II

A single class (class XVI) from the original 2D classification of all autopicked particles was used for a new round of autopicking. This autopick job identified 301,950 particles from the first dataset and 141,167 particles from the second dataset. These 443,117 particles, with a box size of 482×482 Å, were then subjected to 2D classification. 14,592 particles were selected to continue to 3D classification. A preliminary 3D model of this chain was built in Chimera using two heterotrimers interacting at their VPS26 ends; this preliminary model was then filtered to 60 Å and used as input for 3D classification; 6 output classes were used. 13,782 particles were selected for 3D refinement and postprocessing. The final postprocessed chain interface II model had a resolution of 18.6 Å; no B-factor was generated by the postprocessing routine.

### Chain link sub-structure

Processing of the chain link sub-structure centered on VPS26/VPS26 interface began with re-extraction of 15,234 chain interface II particles at a box size of 180×180 Å. Repeated rounds of 2D classification with a mask of 149 Å were performed. 14,275 particles were selected to continue to 3D classification, and a preliminary 3D model generated using only particles from the first dataset was filtered to 60 Å and used as input; 6 output classes were used. 13,392 particles from 3D classification were selected for 3D refinement and postprocessing. The final postprocessed chain link substructure model had a resolution of 6.2 Å and a B-factor of −298.916.

### Tetramer

2D classification of the tetramer particle began with 6,127 particles at a box size of 329×329 Å. 6101 particles continued to 3D classification, which used an initial model generated in cryoSPARC^3^ and filtered to 60 Å; 6 output classes were used. 6015 particles from 3D classification were selected to continue to 3D refinement and posprocessing. The final postprocessed tetramer model had a resolution of 27.4 Å; no B-factor was generated by the postprocessing routine.

